# Cell size is a determinant of stem cell potential during aging

**DOI:** 10.1101/2020.10.27.355388

**Authors:** Jette Lengefeld, Chia-Wei Cheng, Pema Maretich, Marguerite Blair, Hannah Hagen, Melanie R. McReynolds, Emily Sullivan, Kyra Majors, Christina Roberts, Joon Ho Kang, Joachim D. Steiner, Teemu P. Miettinen, Scott R. Manalis, Adam Antebi, Sean J. Morrison, Jacqueline A. Lees, Laurie A. Boyer, Ömer H. Yilmaz, Angelika Amon

## Abstract

Stem cells are remarkably small in size. Whether small size is important for stem cell function is unknown. We find that murine hematopoietic stem cells (HSCs) enlarge under conditions known to decrease stem cell function. This decreased fitness of large HSCs is due to reduced proliferative potential. Preventing HSC enlargement by inhibiting macromolecule biosynthesis or reducing large HSCs size by shortening G_1_ averts the loss of stem cell potential under conditions causing stem cell exhaustion. Finally, we show that a fraction of murine and human HSCs enlarge during aging. Preventing this age-dependent enlargement improves HSC function. We conclude that small cell size is important for stem cell function *in vivo* and propose that stem cell enlargement contributes to their functional decline during aging.

**One Sentence Summary:** Size increase drives stem cell aging.

## MAIN TEXT

Adult stem cells are critical for the maintenance of many tissues in our body. For example, hematopoietic stem cells (HSCs) build the blood system throughout life. Their magnificent ability to proliferate and differentiate into the blood lineages is illustrated by the observation that as few as two HSCs can repopulate the hematopoietic compartment of a lethally irradiated mouse when co-injected with radioprotective bone marrow (BM) (*1*).

As in almost every cell type, HSC division is coupled to cell growth. An increase in cell volume and mass through macromolecular biosynthesis ensures that stem cells retain a constant size after division. Growth control is mediated by the mTOR pathway, which regulates macromolecule biosynthesis in response to amino acid availability and growth signals (*2*). Cell growth ultimately controls cell cycle progression. Cdk4/6-cyclin D complexes drive the decision of HSCs to exit the quiescent state and to divide. They promote cell cycle entry by inactivating the Retinoblastoma (Rb) protein. Rb inhibits the G_1_/S transition by repressing expression of genes required for S phase entry (*3*).

After damage and during aging, stem cell function declines. For example, old HSCs are less competitive in transplantation assays compare to younger ones (*4, 5*). How stem cell fitness declines during aging is only beginning to be understood. In culture, mammalian cells undergo replicative senescence – an irreversible cell cycle arrest (*6–8*). With age, senescent cells accumulate in mice and humans (*9–11*). Whether processes leading to replicative senescence in cultured cells mediate stem cell aging *in vivo* is not clear.

A key characteristic of senescent cells is their large size (*12–14*). Recent work in budding yeast and cultured human cells has provided an explanation for this observation. Senescent cells are large because cell growth and division are only loosely coupled. If cell division is blocked by damage-induced cell cycle checkpoints, macromolecule biosynthesis continues to drive cell growth. As a consequence cells increase in size without a corresponding increase in DNA content. Once the cell cycle arrest has been lifted, cells resume division at a larger size and decreased DNA:cytoplasm ratio (*15–17*). It follows that the more divisions a cell undergoes the more frequently it encounters cell-cycle-arrest inducing damage. Hence, cell size increases and DNA:cytoplasm ratio decreases during replicative aging of yeast and cultured mammalian cells (*12, 18–20*). Importantly, cellular enlargement significantly impacts cell physiology. Young cells manipulated to grow to a large size without a corresponding increase in DNA content exhibit a number of phenotypes observed in senescent cells – foremost, proliferation defects (*16, 21, 22*). Whether large cell size is a cause of senescence and fitness loss during cellular aging *in vivo* is not known.

We show here that cell size affects the function of HSCs *in vivo*. Conditions known to induce stem cell dysfunction - DNA-damage, cell cycle arrest, increased frequency of cell division and aging - cause HSCs to increase in size. Preventing HSC enlargement by interfering with macromolecule biosynthesis or reducing their large size by accelerating progression through G_1_ prevents the loss of stem cell potential. We conclude that cell size is a determinant of stem cell potential and propose that stem cell enlargement contributes to the functional decline of HSCs during DNA-damage and aging.

## RESULTS

### Large cell size contributes to radiation-induced loss of stem cell fitness

Previous studies showed that growing primary human cells to a large size *in vitro* decreases their proliferation potential (*16*). Whether large cell size interferes with proliferation *in vivo* has not yet been determined. We chose HSCs to address this question because of their remarkably small size. To address whether large HSC size is associated with decreased fitness, we first asked whether an insult known to reduce HSC fitness and to induce senescence – DNA damage (*23*) – also causes an increase in HSC size. To induce DNA-damage, we sub-lethally irradiated young mice with 3 gray (Gy). After 2 weeks, we isolated live HSCs (Lin^−^, Sca1/Ly6^+^, CD177/cKit^+^, CD150/Slamf1^+^, CD48/Slamf2^−^, 7-ADD^−^, **Fig. S1A**, **Materials and Methods Fig. 1**) (*1*) and examined their size by Coulter counter and microscopy.

**Fig. 1.**
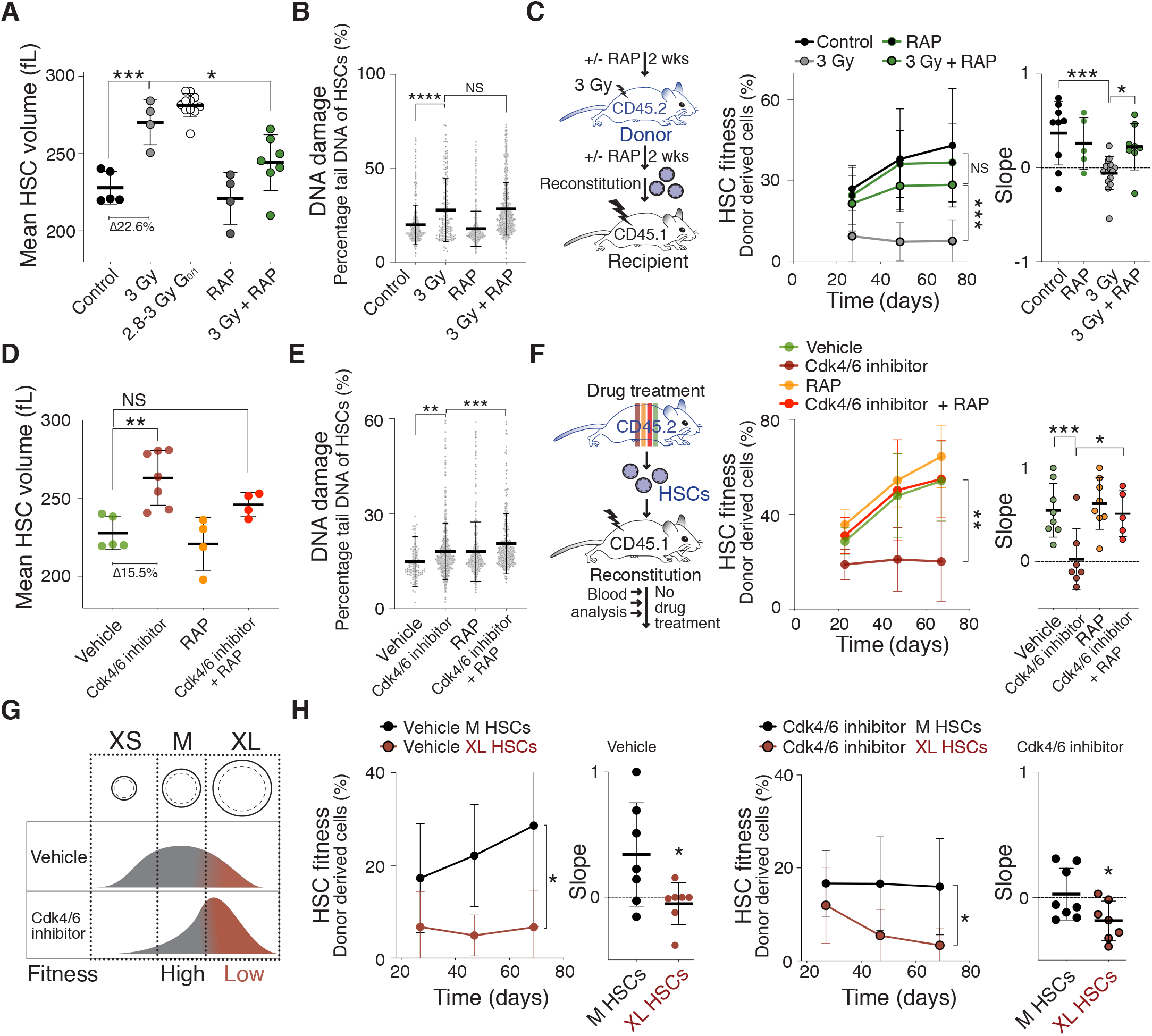
Cellular enlargement contributes to DNA damage-induced fitness decline in HSCs. (**A**) Mean volume (fL) of HSCs obtained from vehicle (*n* = 5), sub-lethally irradiated (3 Gy, *n* = 4), G_0/1_ 2.8-3 Gy (*n* = 11), rapamycin treated (RAP, *n* = 4), and RAP + 3 Gy (*n* = 7) treated mice 2 weeks after irradiation as measured by Coulter counter (Δ = difference). (**B**) Measurement of DNA damage using CometChip: Percentage tail DNA of HSCs (%) isolated from mice 2 weeks after treatment with vehicle, 3 Gy, RAP or 3 Gy + RAP (*n* ≥ 166, 2 independent experiments). (**C**) Reconstitution assay: Donor (CD45.2) mice were pre-treated with rapamycin (RAP) or vehicle for 2 weeks, sub-lethally irradiated (3 Gy) and treated with rapamycin (*n*; donors = 12, recipients = 8) or vehicle (*n*; donors = 12, recipients = 15) for another 2 weeks before 1,000 CD45.2 HSCs were isolated and transplanted together with CD45.1 BM (420,000) into lethally irradiated recipient mice. Control donor HSCs were not treated (control, *n*; donors = 6, recipients = 9) or treated with rapamycin without irradiation (control, *n*; donors = 3, recipients = 5). Recipient mice were not treated with rapamycin after reconstitution. Percentage of donor-derived white blood cells in recipients and slope of reconstitution kinetics over time (3 independent experiments). (**D**) Mean volume (fL) of HSCs isolated from mice treated with vehicle (*n* = 5), Cdk4/6 inhibitor (PD, *n* = 7), rapamycin (RAP, *n* = 4) or Cdk4/6 inhibitor + RAP (*n* = 4) for 85 days was measured using Coulter counter (Δ = difference). Same control as in Fig. 1A. (**E**) Representative measurement of DNA damage in CometChip: Percentage tail DNA of HSCs (%) isolated from mice treated with vehicle, Cdk4/6 inhibitor (PD), RAP or Cdk4/6 inhibitor + RAP (*n* ≥ 552, 3 independent experiments). (**F**) Experimental design to artificially enlarge HSCs: CD45.2 mice were treated with vehicle (*n*; donors = 5, recipients = 8), Cdk4/6 inhibitor (PD, *n*; donors = 5, recipients = 7), rapamycin (RAP, *n*; donors = 5, recipients = 8) or Cdk4/6 inhibitor + RAP (*n*; donors = 5, recipients = 5) for 85 days before their HSCs were isolated for transplantation together with CD45.1 supporting BM (420,000) cells into lethally irradiated CD45.1 recipient mice. No drug treatment was performed after the reconstitution. Percentage (%) of donor-derived white blood cells in recipient and slope of reconstitution kinetics were determined over time (2 independent experiments). (**G**) Experimental strategy to determine the role of cell size for HSC fitness: If size determinates fitness, similarly sized HSCs are expected to exhibit a similar reconstitution potential, irrespective of whether HSCs were treated with vehicle or Cdk4/6 inhibitor (PD). (**H**) Reconstitution assay: M-(*n*; donors = 6, recipients ≥ 7) or XL-(*n*; donors = 6, recipients = 7) sized HSCs of CD45.2 donor-mice treated with vehicle or Cdk4/6 inhibitor (PD) were transplanted together with CD45.1 supporting BM (420,000) cells into lethally-irradiated recipient mice (CD45.1), which were not treated with drugs after the reconstitution. Percentage (%) of donor-derived white blood cells in recipients and slope of reconstitution kinetics were determined over time (3 independent experiments). For all panels, statistical significance was calculated using unpaired *t*-test to compare 2 samples, one-way ANOVA - multiple comparison - Tukey post-hoc test to compare multiple (3 or more) samples, ****P < 0.0001, ***P < 0.001, **P < 0.01, *P < 0.05; NS, not significant. Mean ± s.d is displayed.

With both methods we found that irradiated HSCs were enlarged compared to control HSCs (**Fig. 1A, S1B**). Our analysis further showed that only a fraction of HSCs became enlarged, as judged by a broadening rather than a shift in the size distribution (**fig. S1C**). The size increase of irradiated HSCs was however not only caused by irradiation arresting HSCs in G_2_ phase (*24*). Enlargement was also observed when specifically analyzing G_0/1_ HSCs (**Fig. 1A**).

To determine whether large HSC size was associated with irradiation-induced senescence, we correlated cell size with senescence-associated beta-galactosidase (SA-β-gal) 2 weeks after either 3 Gy or 6 Gy irradiation (*23*). We did not detect high SA-β-gal activity in HSCs obtained from 3 Gy-irradiated or control animals (**fig. S1D-E**). In contrast, SA-β-gal^high^ HSCs were readily observed in 6 Gy-irradiated animals. Next, we sorted HSCs based on SA-β-gal activity levels and found that SA-β-gal^high^ HSCs were larger than SA-β-gal^low^ HSCs from the same animal (**fig. S1F-H**). Thus, low doses of irradiation (3 Gy) cause HSC enlargement without detectable induction of SA-β-gal activity, whereas a higher dose (6 Gy) leads to enlarged HSCs that harbor high levels of SA-β-gal activity. These observations indicate that limited DNA damage enlarges HSCs before inducing a bona-fide senescence program and that large HSCs are more likely to be senescent.

To determine whether increased HSC size contributes to their irradiation-induced dysfunction, we prevented HSC enlargement during irradiation. We treated mice with rapamycin (mTOR inhibitor) or vehicle for two weeks, irradiated them with 3 Gy or left the mice untreated (control), followed by a rapamycin or vehicle treatment for another 2 weeks. Rapamycin treatment prevented HSCs from increasing in size in response to irradiation (3 Gy + RAP, **Fig. 1A, S1B**), without changing the cell cycle state of HSCs as measured by DNA content and EdU incorporation (**fig. S2A-B**). Importantly, rapamycin treatment neither decreased the level of DNA damage after 3 Gy irradiation as evaluated by alkaline comet assays (**Fig. 1B, S2C**), nor altered the differentiation potential of HSCs as demonstrated by blood lineage analysis after reconstitution (**fig. S2D**). We conclude that rapamycin prevents HSC enlargement after irradiation.

Having established that rapamycin prevents HSC enlargement without affecting the degree of DNA damage, we assessed the effects of rapamycin treatment on HSC function. We isolated HSCs from donor mice expressing the pan-hematopoietic lineage allele CD45.2 that were treated with rapamycin or vehicle during 3 Gy irradiation. We then lethally irradiated recipient mice carrying the CD45.1 allele to eliminate the pre-existing blood system. The recipient mice were reconstituted with CD45.2 donor HSCs and supporting CD45.1 bone marrow cells to allow for donor-driven reconstruction of the blood system (fitness). Donor HSC fitness was determined by the fraction of peripheral CD45.2 white blood cells in CD45.1 recipients over time. Rapamycin had a dramatic effect on the fitness of irradiated HSCs: Donor HSCs from mice that were 3 Gy irradiated and received rapamycin treatment exhibited a significantly increased reconstitution potential compared to HSCs from 3 Gy irradiated mice (**Fig. 1C, S2E**). Importantly, the degree of chimerism changed over time. The contribution to the blood system of 3 Gy irradiated HSCs treated with rapamycin increased over the course of the reconstitution, whereas the fitness of vehicle-treated 3 Gy irradiated HSCs decreased (**Fig. 1C**; slope analysis). This observation indicates that HSC proliferation, rather than homing to the bone marrow, is affected by rapamycin. We note that rapamycin was not universally radioprotective. The drug failed to confer radioprotection at higher radiation doses (5 Gy; **fig. S2F**) despite suppressing SA-β-gal activity levels (**fig. S1E**). We conclude that rapamycin prevents radiation-induced HSC exhaustion. We interpret this result to suggest that when cellular enlargement is prevented by rapamycin during DNA damage, HSCs retain their fitness.

### The Cdk4/6 inhibitor palbociclib enlarges HSCs causing their decline in reconstitution potential

How does irradiation lead to HSC enlargement? Directly after irradiation, HSCs experience DNA damage and arrest in G_2_ to repair this damage (*24*). This arrest is, however, transient. HSC populations return to their pre-irradiation cell cycle state 2 weeks after sub-lethal irradiation (**fig. S2A-B**). Despite the return of HSCs to the G_0/1_ state, their size increased (**Fig. 1A**). *In vitro* studies with budding yeast and primary human cells provide a potential explanation for this finding. During cell cycle arrest, mTOR continues to promote macromolecule biosynthesis and cells increase in size (*15–17*).

To determine whether cell cycle arrest causes HSCs to enlarge, we examined HSC size in response to treating animals with the Cdk4/6 inhibitor palbociclib (PD), which arrests cells in G_1_ phase (*25*). Because HSCs rarely divide under physiological conditions (*26*), we injected PD every other day for 120 days (**fig. S3A**). This treatment regimen increased G_0/1_ length as judged by cell cycle analysis and changes in blood composition of donor animals (**fig. S3B-E**). Analysis of HSC size after various lengths of PD treatment showed that HSCs increased in size in a time-dependent manner (**fig. S3A**). After 85 days of PD treatment, the mean volume of HSCs was 263 fL as measured by Coulter counter, which reflects an increase in size of 15.5 % compared to control HSCs (**Fig. 1D**). These data indicate that delaying cells in G_0/1_ by PD treatment causes HSC enlargement *in vivo.*

Characterization of PD-enlarged HSCs revealed increased DNA damage (**Fig. 1E**), which was not associated with the production of SA-β-gal activity (**fig. S3F**). The finding that a PD-arrest is associated with DNA damage, causal or consequential, afforded us another opportunity to assess whether large HSCs that harbor DNA damage exhibit decreased reconstitution potential. A previous study showed that short-term inhibition of Cdk4/6 (12 h, trilaciclib) did not reduce HSC reconstitution potential (*25*). Cell size, however, is not expected to increase within the treatment time frame of this study. We found that prolonged PD treatment enlarged HSCs and impaired their reconstitution potential (**Fig. 1F, S3G**). To determine whether the large size of PD-treated HSCs contributed to their reduced reconstitution potential, we prevented size increase during PD treatment by simultaneously treating animals with rapamycin. When treated with PD *and* rapamycin, HSCs DNA damage levels were not affected but HSCs did not increase in size (**Fig, 1D, E**). Importantly, their reconstitution potential was preserved (**Fig. 1F**). Thus, rapamycin prevents PD-induced enlargement of HSCs and preserves their fitness.

We hypothesized that if PD treatment indeed decreases fitness because it enlarges HSC size, then HSCs of PD-treated animals that did not grow to a large size should exhibit a reconstitution potential similar to that of untreated animals. In other words, medium (M)-sized HSCs from PD-treated mice should exhibit a greater reconstitution potential than large (XL)-sized HSCs from PD- or vehicle-treated mice (**Fig. 1G)**. We found this to be the case. To obtain M-sized and XL-sized HSCs from PD-arrested or vehicle-treated mice, we used the forward scatter during cytometric sorting. We isolated the largest 10 % of HSCs (XL) and HSCs of a mean size +/− 10 % (M) (*27*). Cell volume analysis confirmed that we isolated PD- or vehicle-treated HSCs of the same absolute size using the M-(246 fL) or XL-gate (305 fL, 24 % larger, **fig. S3H**). M-sized PD-treated HSCs were more effective in reconstituting lethally irradiated mice than XL-sized PD- or vehicle-treated HSCs (**Fig. 1H**). These results indicate that PD-induced enlargement rather than PD-treatment *per se* causes HSCs to be less fit. We note that M-size PD-treated HSCs appear less fit than vehicle-treated, M-sized HSCs when comparing reconstitution kinetics (slope). This difference is however not statistically significant. It is nevertheless possible that PD treatment also affects HSC potential in a cell size independent manner. We conclude that PD reduces HSC fitness largely by increasing their cell size.

### Cellular growth enlarges HSCs and reduces their reconstitution potential

Large size contributes to the decreased reconstitution potential of HSCs with DNA damage. To determine whether enlargement alone decreases HSC fitness *in vivo*, we analyzed HSCs that were large but lacked DNA damage. To create large undamaged HSCs, we manipulated mTOR pathway activity. The lack of *TSC1* causes constitutive activation of mTOR signaling, which increases cell growth and enlarges the size of blood cells (*28*). Cre-driven excision of *TSC1* in *TSC1^fl/fl^;R26-creERT2* mice (henceforth *TSC1^−/−^*) enlarged HSCs by 27-60 % without causing increased DNA damage (**Fig. 2A-B**). Next, we isolated HSCs from *TSC1^fl/fl^;R26-creERT2* mice and reconstituted lethally irradiated recipients (**Fig. 2C**). We allowed HSCs to home to the bone marrow in recipients for 3 days, then induced *TSC1* excision using tamoxifen and assessed donor *TSC1^−/−^* HSC fitness. *TSC1*^−/−^ HSCs were impaired in repopulating the blood system of recipient mice (**Fig. 2D**). We conclude that constitutive mTOR-driven growth enlarges HSCs without causing DNA damage, yet reduces their ability to reconstitute lethally irradiated recipients. These data support the conclusion that increased cell growth enlarges HSCs and decreases their fitness.

**Fig. 2.**
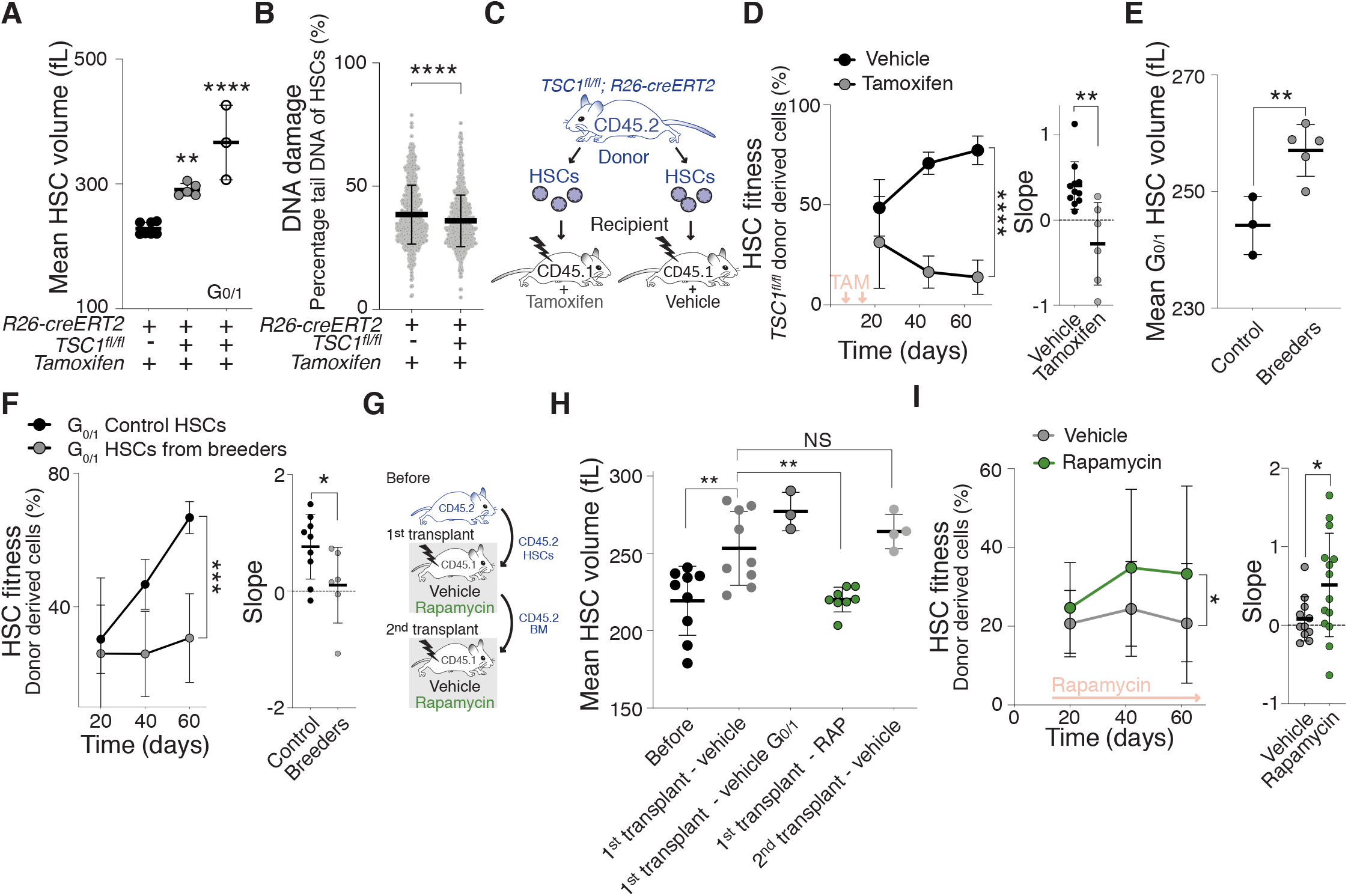
mTOR hyperactivation and high cell division frequency enlarge HSCs contributing to their fitness decline. (**A**) Mean volume (fL) of *TSC1*^+/+^ (*n* = 5), *TSC1*^−/−^ (*n* = 5) and G_0/1_ *TSC1*^−/−^ HSCs (*n* = 3) 60 days after vehicle or tamoxifen treatment was measured using a Coulter counter. Same control as in Fig. 1A. (**B**) CometChip assay to measure DNA damage: Percentage tail DNA of HSCs (%) from *TSC1*^+/+^ (*n* = 769) or *TSC1*^−/−^ (*n* = 838) mice. (**C**) 800 HSCs from *TSC1^fl/fl^;R26-creERT2* mice were transplanted together with CD45.1 supporting BM (420,000) cells into lethally irradiated recipient mice and, were allowed to home to the bone and the recipients were then treated with vehicle or tamoxifen to induced *TSC1* excision 3 and 14 days after transplantation. (**D**) Reconstitution assay described in (C): Percentage (%) of *TSC1*^+/+^ (*n;* donors = 7, recipients = 11) and *TSC1*^−/−^ (*n;* donors = 7, recipients = 7) donor HSC-derived white blood cells in recipients and slope of reconstitution kinetics were determined over time. Red arrows indicate recipient treatment with tamoxifen or vehicle (3 independent experiments). (**E**) Mean volume (fL) of G_0/1_ HSCs obtained from age-matched virgin (*n* = 3) and breeding (*n* = 5) female mice was measured using a Coulter counter. (**F**) Reconstitution assay: 600 G_0/1_ HSCs were isolated from age-matched virgin (*n;* donors = 6, recipients = 9) or breeding (*n;* donors = 5, recipients = 6) donor (CD45.2) mice and transplanted together with CD45.1 BM (420,000) into lethally irradiated recipient mice. Percentage (%) of donor-derived white blood cells in recipients and slope of reconstitution kinetics were determined over time. (**G**) Scheme of transplantation experiment (H-I): Following transplantation, animals were treated with rapamycin or vehicle for 80 days. HSC size was analyzed before transplantation and 80 days after the 1^st^ and 2^nd^ transplantation. BM from 1^st^ transplant was used for 2^nd^ transplant. (**H**) Mean volume (fL) of all CD45.2 HSCs before transplantation (*n* = 9) and 80 days after treatment with vehicle (*n* = 9), vehicle G_0/1_ (*n* = 3) or rapamycin (RAP, *n* = 8) treatment during 1st or 2^nd^ transplantation (*n* = 4) was measured using a Coulter counter. (**I**) Reconstitution assay: Donor (CD45.2)-derived HSCs together with supporting BM (CD45.1) (420,000) were transplanted into lethally irradiated recipient mice that were treated with vehicle (*n;* donors = 10, recipients = 11) or rapamycin (*n;* donors = 10, recipients = 14) for 80 days following reconstitution. Percentage (%) of donor-derived white blood cells in recipients and slope of reconstitution kinetics were determined over time (3 independent experiments). For all panels, statistical significance was calculated using unpaired *t*-test to compare 2 samples, one-way ANOVA - multiple comparison - Tukey post-hoc test to compare multiple (3 or more) samples, ****P < 0.0001, ***P < 0.001, **P < 0.01, *P < 0.05; NS, not significant. Mean ± s.d is displayed.

### Consecutive cell divisions cause HSCs to enlarge, which drives their exhaustion

We hypothesized that if cellular enlargement contributes to HSC dysfunction, any condition causing loss of stem cell potential ought to be accompanied by HSC enlargement. Successive cell divisions cause stem cell exhaustion. With each cell division, HSCs lose their cell division potential, such that each daughter is less potent than its mother (*29–31*). We determined whether HSCs enlarge during two conditions known to cause higher levels of cell divisions *in vivo*: pregnancy (*32*) and transplantation into irradiated mice (*4*). G_0/1_ HSCs of breeding females and HSCs obtained after transplantation were enlarged (**Fig. 2E, G-H**). As in response to radiation, not all HSCs enlarged during serial transplantation, as judged by a broadening of the size distribution (**fig. S4A**). This was not due to changes in cell cycle distribution. 80 days after transplantation (the time cell size was analyzed), cell cycle distribution of HSCs had returned to the same cell cycle profile as before transplantation (**fig. S4B-C**). Interestingly, in secondary transplants, donor HSCs did not increase further in size (**Fig. 2H)**, suggesting that an upper size threshold exists for HSCs.

The two methods of increasing the rate of HSC division also caused a decrease in reconstitution potential (**Fig. 2F, S4D**). This decreased fitness was not due to transplanting actively proliferating HSCs (*33*), because we only transplanted HSCs in G_0/1_. We conclude that similar to budding yeast and primary mammalian cell *in vitro* (*12, 18*), successive divisions of HSCs are accompanied by an increase in HSC size and decrease in fitness.

The reconstitution experiment afforded us the opportunity to determine whether cell division-driven enlargement promotes HSC exhaustion (*31*). We conducted transplantation experiments, in which <u>recipients</u> received prolonged rapamycin treatment to prevent HSC enlargement during the many divisions needed to reconstitute the hematopoietic compartment (**Fig. 2G**). This experimental regime indeed prevented donor HSCs from enlarging (**Fig. 2H**). Importantly, preventing enlargement of donor HSCs in the host via rapamycin treatment improved their fitness during the reconstitution (**Fig. 2I**). We conclude that successive divisions cause HSCs to enlarge, which reduces their stem cell potential.

Why does an increased cell division frequency cause HSC enlargement? Stochastic damage can occur during every cell division (*34, 35*). When cells are damaged, cell cycle checkpoints are activated to facilitate repair (*24, 36*). During this arrest, mTOR continues to promote macromolecule biosynthesis driving cell growth and enlargement (*15, 16, 21, 22*). It follows that the more divisions a cell undergoes, the more likely it encounters damage that causes cell-cycle-arrest and therefore enlargement. Our data indicate that in HSCs too successive divisions cause enlargement, which in turn leads to HSC dysfunction. We note that this hypothesis also explains why only a fraction of HSCs enlarge during serial transplantations.

### Naturally large HSCs display decreased reconstitution potential

Our data show that DNA damage, proliferation challenge or artificially increasing cell size decreases HSC fitness. To determine the broader physiological relevance of this observation, we assessed the fitness of naturally large HSCs. We isolated the smallest 10 % of HSCs (XS), the largest 10 % of HSCs (XL) and HSCs of a mean size +/− 10 % (M; **Fig. 3A**) from young wild-type mice by flow cytometry using forward scatter. Cell volume analysis by Coulter counter confirmed that this method is able to differentiate HSCs based on their size when size differences are large (**Fig. 3B-C, Materials and Methods Fig. 1**). XS HSCs (mean = 209 fL) were 30% smaller than M HSCs (mean = 240 fL), which were 34% smaller than XL HSCs (mean = 312 fL).

**Fig. 3.**
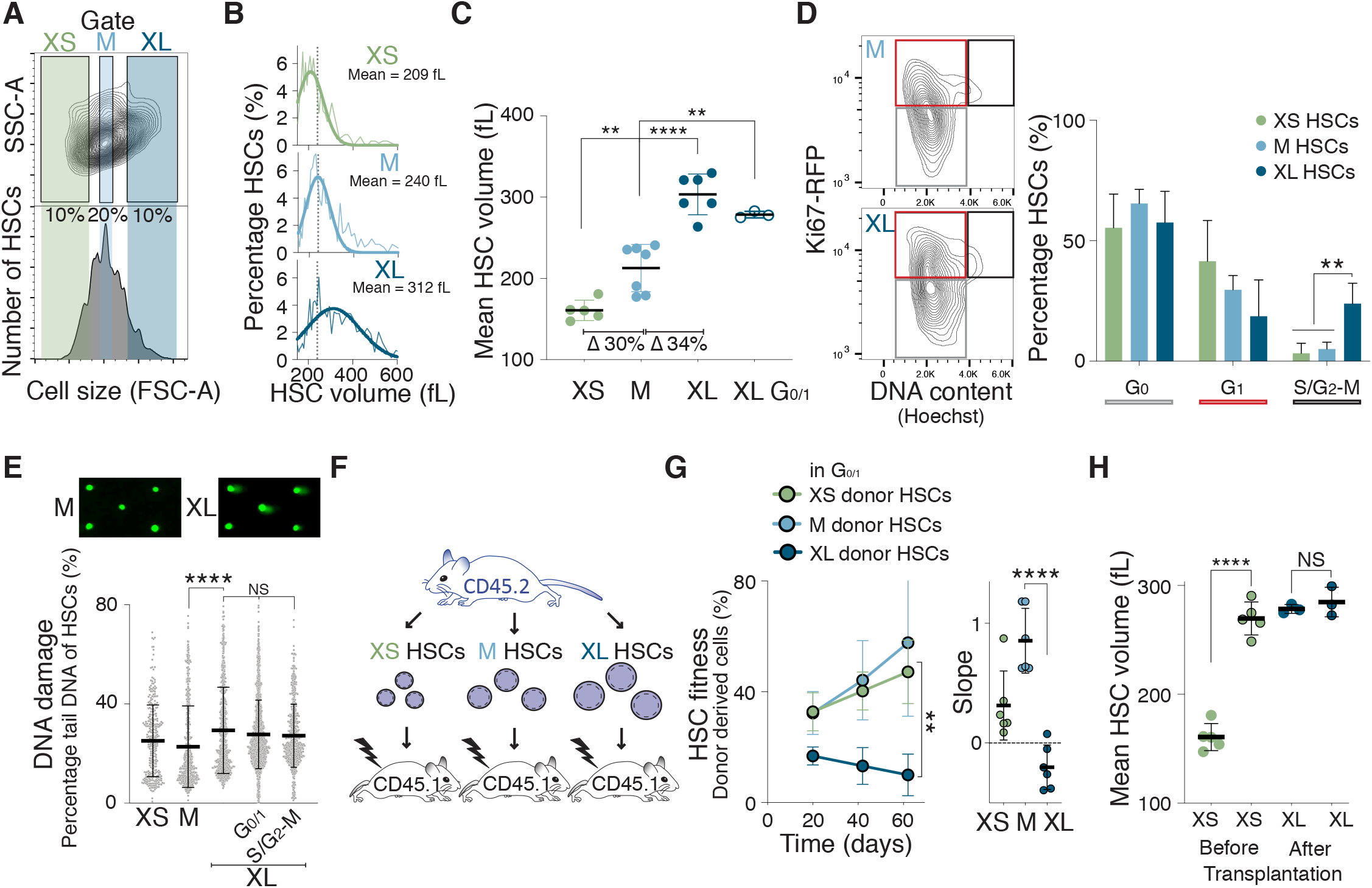
Naturally large HSCs are impaired in reconstituting the hematopoietic compartment. (**A**) Size distribution of HSCs as determined by forward scatter (FSC-A). Gates used to isolate small (XS), medium (M) and large (XL) HSCs are indicated. (**B**) Percentage of HSCs per volume (fL) isolated using the XS, M or XL gates shown in (A) measured by Coulter counter. Gaussian fit was used to determine mean cell volume. Dotted line marks the mean of M-sized HSCs in all images. (**C**) Mean volume (fL) of HSCs isolated using the XS (*n* = 5), M (*n* = 7), XL (*n* = 6) or G_0/1_ XL (*n* = 3) gates (Δ = difference) was measured using a Coulter counter. (**D**) Representative Ki67-RFP and DNA content analysis. The percentage of XS-, M- or XL-HSCs in the G_0_, G_1_ and S/G_2_-M phases of the cell cycle was determined by DNA content (Hoechst 33342) and Ki67-RFP analysis (*n* = 4). (**E**) CometChip assay to measure DNA damage: Percentage tail DNA of XS-, M- and XL, G_0/1_ XL and S/G_2_-M XL-sized HSCs (*n* = 2 independent experiments with > 323 cells measured per condition). (**F**) Schematic of reconstitution experiments: 600 differently sized CD45.2 (blue) donor-derived HSCs together with CD45.1 supporting BM cells (420,000) were transplanted into lethally irradiated recipient mice. (**G**) Reconstitution assay: Percentage (%) of donor-derived (CD45.2) white blood cells in recipients after transplantation of G_0/1_ Hoechst-labeled donor XS-HSCs (*n;* donors = 5, recipients = 6), M-HSCs (*n;* donors = 5, recipients = 6) or XL-HSCs (; donors = 5, recipients *n* = 6) shown in (F) over time (2 independent experiments). Slope of reconstitution kinetics over time is also shown. (**H**) Mean volume (fL) of XS- or G_0/1_ XL-sized CD45.2 HSCs before transplantation and G_0/1_ XS- or G_0/1_ XL-sized CD45.2 HSCs 80 days after transplantation as measured by Coulter counter (*n* ≥ 3). Before data as in Fig. 3C and after as in Fig. 5G. For all panels, statistical significance was calculated using unpaired *t*-test to compare 2 samples, one-way ANOVA - multiple comparison - Tukey post-hoc test to compare multiple (3 or more) samples, ****P < 0.0001, ***P < 0.001, **P < 0.01, *P < 0.05; NS, not significant. Mean ± s.d is displayed.

We next investigated why XL HSCs are large. During cell division, cells increase in size to maintain constant volumes of the two daughter cells. To determine whether XL HSCs were large because they were in the process of dividing, we analyzed their DNA content. While the vast majority of differently sized HSCs were in G_0,_ XL-HSCs were less in G_1_ and more often in S phase or mitosis (S/G_2_-M, **Fig. 3D, S5A-B**). As XL HSCs are not polyploid (*37*), these data indicate that the XL HSC population is comprised of large G_0/1_ HSCs and HSCs that are larger because they are in S/G_2_-M.

To determine whether naturally large G_0/1_ HSCs harbor DNA damage, we performed an alkaline comet-chip assay. This analysis revealed that G_0/1_ XL HSC nuclei displayed a higher percentage of tail DNA compared to XS- and M-sized HSCs and higher levels of γH2AX phosphorylation (**Fig. 3E, S5C**). We conclude that WT mice harbor large G_0/1_ HSCs, which have more DNA damage compared to small WT HSCs.

To assess the fitness of naturally XL HSCs *in vivo*, we isolated XS, M and XL HSCs and reconstituted lethally irradiated recipient mice (**Fig. 3F**). Because HSCs in S/G_2_-M are impaired in providing long-term multi-lineage reconstitution (*33, 38*), we specifically isolated G_0/1_ HSCs using a DNA stain. Remarkably, the reconstitution potential differed with HSC size. XS- and M-sized G_0/1_ HSCs effectively contributed to the peripheral blood of recipient mice, whereas XL G_0/1_ HSCs exhibited a significantly reduced reconstitution potential (**Fig. 3G, S5D, F**). DNA staining can alter HSCs fitness (*39*), but reconstitutions without staining the DNA of HSCs revealed similar results (**fig. S5E**). XS HSCs enlarged during reconstitutions, while G_0/1_ XL HSCs remained large by the end of the reconstitution experiment (**Fig. 3H**). This indicates that XL HSCs do not shrink back to their original size during the divisions necessary to rebuild the recipients blood system or that they are incapable of dividing altogether.

In summary, we have examined six different experimentally induced or physiological conditions, under which HSCs exhaustion occurs: irradiation-induced DNA-damage, Cdk4/6 inhibition, hyperactivation of mTOR, high division rate during transplantation, repeated pregnancy, and naturally large HSCs. We observe that HSC enlarge under all these conditions. Importantly, in all experimental settings, in which this could be investigated, preventing HSC enlargement with rapamycin preserved HSC fitness.

### Large HSCs exhibit decreased proliferation potential

Why are large HSCs less fit than small HSCs? Large size could affect (1) proliferation potential, (2) stem cell identity and differentiation potential, (3) the metabolic state of HSCs and/or (4) protein synthesis capacity. We chose naturally large and PD-enlarged HSCs to test these possibilities.

#### (1) Proliferation potential

The observation that PD-enlarged and XL-donor HSCs contribute less to the blood lineages of recipient animals over time (reconstitution slopes) indicated that cell size affects HSC proliferation. The analysis of HSC proliferation *in vitro* confirmed this conclusion. We found that PD-enlarged HSCs formed fewer colonies than control HSCs (**Fig. 4A**). Rapamycin treatment partially restored colony formation of PD-treated HSCs. A decrease in proliferation was also observed in naturally large HSCs, although the effect was less striking (**Fig. 4A**). We also investigated whether large size affected homing of HSCs to the bone marrow niche, which is essential for hematopoiesis. We injected XL or PD-enlarged donor HSCs into CD45.1 recipient mice and examined the fraction of CD45.2 donor cells in the bone marrow 21 hours thereafter. This analysis suggested that HSC size did not affect homing potential (**fig. S6A**). We conclude that cellular enlargement interferes with the ability of HSCs to proliferate.

**Fig. 4.**
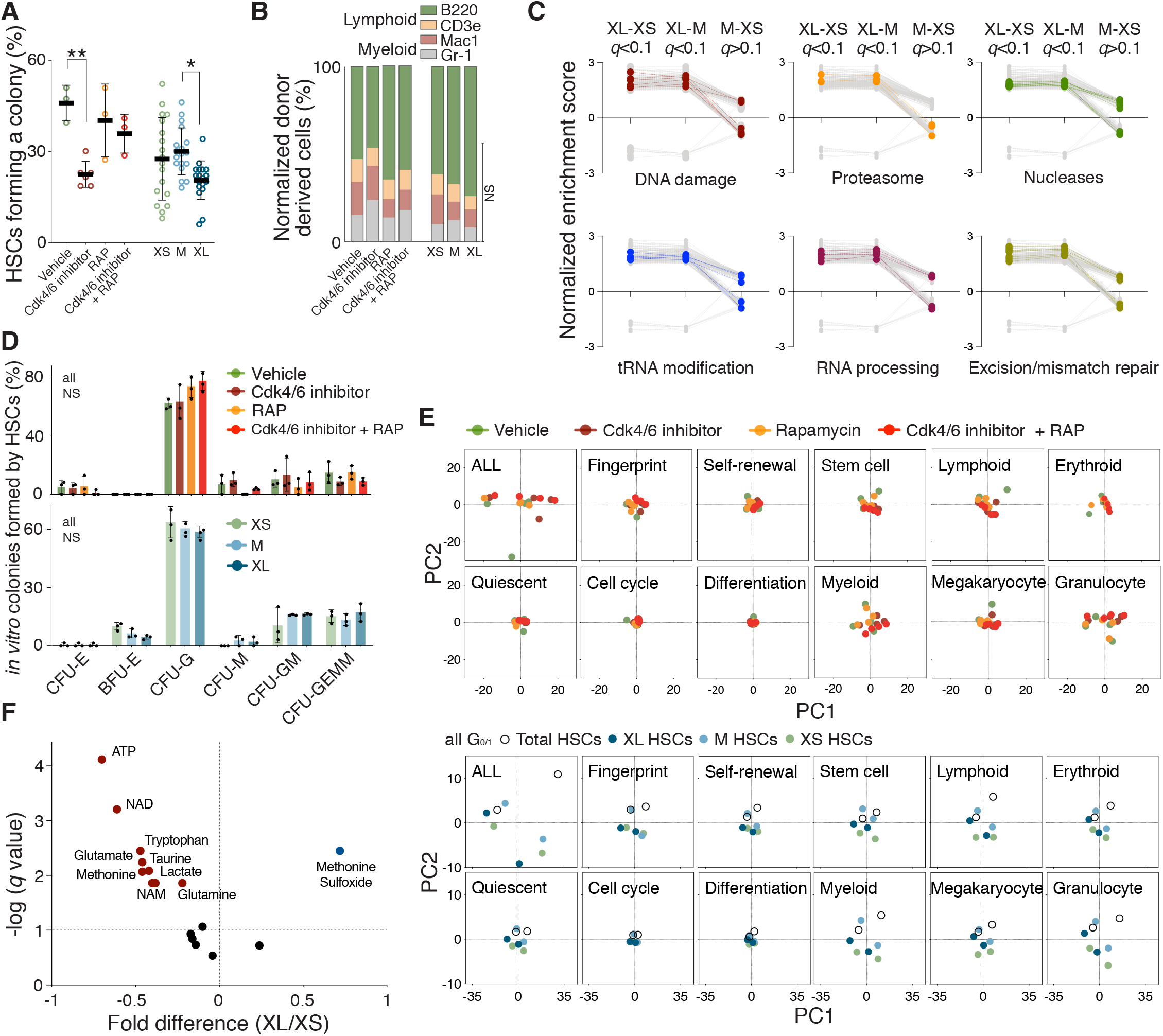
Large hematopoietic stem cells exhibit reduced proliferation capacity. (**A**) Colony forming efficiency *in vitro*: HSCs (w/ and w/o Hoechst staining) were isolated from mice treated with vehicle, Cdk4/6 inhibitor (PD), rapamycin (RAP) or Cdk4/6 inhibitor + RAP (*n* ≥ 3) and XS-, M- or XL-HSCs (*n* = 18) from untreated WT mice. HSCs were plated onto methylcellulose and the percentage of HSCs forming a colony was quantified. HSCs were not treated with drugs *ex vivo*. (**B**) *In vivo* differentiation assay: Relative lineage distribution (lymphoid lineage B220, CD3e; myeloid lineage Mac1, Gr-1) from peripheral donor-derived (CD45.2) white blood cells 80 days after recipient mice were reconstituted with HSCs from mice treated with vehicle (*n* = 8), Cdk4/6 inhibitor (PD, *n* = 6), RAP (*n* = 8) or Cdk4/6 inhibitor + RAP (*n* = 5) or with XS-(*n* = 5), M-(*n* = 12), XL-(*n* = 6) HSCs from WT mice. No drug treatment was performed after the reconstitution. (**C**) GO gene sets were analyzed using GSEA and then filtered to be different (*q* < 0.1) when comparing M- and XS-HSCs with XL-HSCs (all G_0/1_), but indistinguishable (*q* > 0.1) when comparing M-HSCs with XS-HSCs. Biological processes are colored when ≥ 20 % of their GO gene sets passed the filter. Biological processes are displayed in grey when less than 20 % of their GO gene sets passed the filter (**Table S1**). (**D**) *In vitro* differentiation assay: HSCs from mice treated with vehicle, Cdk4/6 inhibitor (PD), rapamycin (RAP) or Cdk4/6 inhibitor + RAP (top graph) or XS-, M- or XL-HSCs (bottom graph) were plated onto methylcellulose and GEMM, GM or single lineages (G, M and E) colonies were counted (*n* = 2 independent experiments with 3 replicates each, representative experiment shown). (**E**) Expression differences between differently sized G_0/1_ HSCs were assessed on the basis of principle components analysis (PCA) using selected HSC genes affecting stem cell identity and differentiation (**Table S2**). Relative distances were collapsed into two dimensions using PCA. Samples analyzed: total, XS-, M-, XL HSCs (*n* = 2, bottom) and vehicle, Cdk4/6 inhibitor (PD), rapamycin (RAP), Cdk4/6 inhibitor + RAP treated HSCs (*n* = 4, top). (**F**) Volcano plot representing LC-MS measurements for metabolites in G_0/1_ XL-HSCs compared to G_0/1_ XS-HSCs. The x-axis represents the fold-change and the y-axis represents adjusted FDR. Colored dots represent metabolites that change with size significantly (*n* = 7 independent experiments). For all panels, statistical significance was calculated using unpaired *t*-test to compare 2 samples, one-way ANOVA - multiple comparison - Tukey post-hoc test to compare multiple (3 or more) samples, ****P < 0.0001, ***P < 0.001, **P < 0.01, *P < 0.05; NS, not significant. Mean ± s.d is displayed.

#### (2) Stem cell identity and differentiation potential

To determine whether cell size affects HSC identity or their ability to differentiate into the various hematopoietic lineages, we first conducted a comparison of gene expression profiles between differently sized HSCs using Gene Set Enrichment Analysis (GSEA). This analysis revealed that the expression pattern of several GO gene sets changed in G_0/1_ XL HSCs compared to smaller control HSCs (M and XS). To identify transcriptional changes in XL-HSCs that underlie the decline in fitness *in vivo*, we focused our analysis on gene ontology (GO) sets that specifically changed in XL-HSCs relative to XS- and M-HSCs, but not between XS- and M-HSCs. 2.4% of all GO gene sets passed this filter (**Table S1**). Using a cut-off that required that > 20 % of GO gene sets associated with a given biological process to be differentially regulated between XL HSCs versus XS and M HSCs, we identified the following pathways (**Fig. 4C**): DNA damage (21 % of associated GO gene sets were differentially regulated in XL vs XS/M sized HSCs), proteasome (30 %), excision/mismatch repair (50 %), RNA processing (32 %), tRNA modification (23 %), and nucleases (27 %). Importantly, the up-regulation of genes involved in DNA damage response and repair was biologically relevant. Naturally large HSCs harbored more DNA damage than smaller control HSCs (Fig. 3E). In the comparison between PD enlarged HSCs to control HSCs only 0.3% of the GO gene sets passed the functional filter and no enrichment for particular biological processes was observed (**Table S1**).

To specifically determine whether size affected the self-renewal or differentiation ability of HSCs, we used published lists of genes related to HSC identity and differentiation (**Table S2**) and analyzed whether their expression changed in XL and PD-enlarged HSCs. We found that only a few genes rose to the level of significant differential expression (**Table S3**). A principle component analysis confirmed that size had little or no effect on the expression of these gene categories (**Fig. 4E**). To evaluate the biological relevance of this finding, we employed functional assays to assess differentiation potential of large sized HSCs *in vitro* and *in vivo*. The ability to form cells of the myeloid, B-, and T-cell lineages did not significantly differ between mice reconstituted with differently sized HSCs (**Fig. 4B**). Differentiation was also not significantly affected *in vitro*. The types of colonies formed by HSCs in methylcellulose cultures were similar between HSCs of different sizes (**Fig. 4D**). Lastly, the concentration of stem cell surface markers was similar between differently sized HSCs (**fig. S6B**). We conclude that cell size does not significantly affect stem cell identity or differentiation potential of HSCs.

#### (3) Metabolic state

A prior study suggested that muscle stem cells can enter a metabolically activated state, termed G_alert_, which enables rapid cell cycle commitment in response to stress. G_alert_ is characterized by an increase in cell size, high mitochondrial activity, high ATP levels and upregulation of the INF-γ response (*40*). To determine whether HSC enlargement could reflect HSCs in such a G_alert_ state, we queried characteristics of the G_alert_ state in large HSCs. In contrast to G_alert_ muscle stem cells, mitochondrial function was not increased in large HSCs as judged by mitochondrial concentration (**fig. S6C)**, reactive oxygen species concentration (**fig. S6D**), expression of mitochondrial genes (**fig. S6E**) and quantification of the mitochondrial transcription factor Tfam (**fig. S6F**). Increased Tom20 staining potentially suggests that mitochondrial protein import is increased in XL-HSCs (**fig. S6F**). Naturally large HSCs were also not metabolically activated. Liquid chromatography-mass spectrometry revealed that G_0/1_ XL-HSCs harbored significantly lower levels of ATP than XS-sized HSCs (**Fig. 4F**). Finally, we found that 40 % of GO gene sets associated with the INF-response were lower in XL-HSCs compared to smaller HSCs, but not significantly different between smaller HSCs (XS and M) using a relaxed functional filter (*q*-val = 0.5, **fig. S6G**). We conclude that naturally large HSCs do not exhibit characteristics of the G_alert_ state.

#### (4) Protein synthesis capacity

Enlarged cultured primary human cells experience decreased protein synthesis rates and ribosome loss that leads to cytoplasm dilution (*16, 41*). To assess protein synthesis capacity in naturally large HSCs, we measured nucleolar size as this often correlates with rRNA synthesis and ribosome biogenesis (*42, 43*). Nucleolar size scaled with HSC size (**fig. S6H**) suggesting that ribosome biogenesis does not decline in large HSCs. Consistent with this conclusion, cell density, to which ribosomes are a significant contributor (*41*), was not affected in large HSCs either (**fig. S6I**). In summary, our characterization of naturally large and PD-enlarged HSCs indicates that large size primarily interferes with the ability of HSCs to proliferate.

### Large size decreases the colony-forming potential of intestinal stem cells

Does cell size also affect the proliferation potential of other adult stem cell types? To address this question, we used Lgr5-GFP knock-in mice to isolate intestinal stem cells (ISCs), which can be distinguished from transient amplifying cells by a two-fold or higher intensity of Lgr5-GFP (**fig. S7A**). We isolated the smallest 15 % of ISCs, the largest 15 % and the mean-sized population +/− 15 % using the forward scatter as a measure of cell size (**fig. S7B**). Cell volume measurements confirmed that this method distinguished ISCs by their size (**fig. S7C**). Furthermore, like naturally large HSCs, XL-ISCs harbored more DNA damage than smaller ones (**fig. S7D**). They also shared other characteristics with XL HSCs. They exhibited a reduced ability to form intestinal organoids (**fig. S7E**), a colony formation assay for ISCs. Importantly, among all the differently sized ISCs, XL ISCs displayed the highest intensity of Lgr5-GFP (**fig. S7F**), suggesting that the colony-forming defect observed in XL-ISCs was not due to them having differentiated into Lgr5^low^ progenitors. We conclude that, like in HSCs, DNA damage is associated with an increase in ISC size and a decrease in colony-forming potential.

### Reducing the size of large HSCs restores their reconstitution potential

Our results indicate that preventing HSC enlargement with rapamycin during DNA-damage or during high frequency of cell division preserves HSC fitness. Although, rapamycin’s main function is to inhibit mTOR-mediated macromolecule biosynthesis, it was nevertheless possible that rapamycin protected HSCs by other mechanisms. To address this possibility, we modulated HSC size by means other than rapamycin treatment. We shortened G_1_.

Inactivation of the S-phase entry inhibitor Rb accelerates progression through G_1_ and reduces cell size (*44*). We hypothesized that reducing the size of large HSCs by reducing Rb levels, improves HSC fitness. To reduce Rb levels in HSCs, we utilized mice that were heterozygous for an excisable *RB* allele (*RB^fl/+^;R26-creERT2*). We first sub-lethally irradiated mice and allowed HSCs to enlarge for 7 days (**Fig. 5A-B**). We then excised one *RB* allele by administering tamoxifen for another 7 days. Loss of one copy of *RB* indeed led to decreased cell size after irradiation (**Fig. 5B**). To determine whether reducing the size of previously enlarged HSCs improved their reconstitution potential, we transplanted these irradiated HSCs into recipients (**Fig. 5C**). This analysis revealed that the lack of one *RB* copy affected the fitness of previously irradiated, large HSCs: Irradiated *RB*^+/−^ HSCs built 15 % of the blood cells of reconstituted animals, whereas irradiated *RB*^+/+^ HSCs contributed only 5 %, which is consistent with our hypothesis that DNA-damaged and *RB* deficient HSCs divide more (**fig. 87A**) (*45*). Importantly, during the reconstitution, these irradiated *RB*^+/−^ HSCs were smaller in size than irradiated *RB*^+/+^ HSCs (**Fig. 5D**). We conclude that reducing cell size improves the reconstitution potential of irradiated HSCs.

**Fig. 5.**
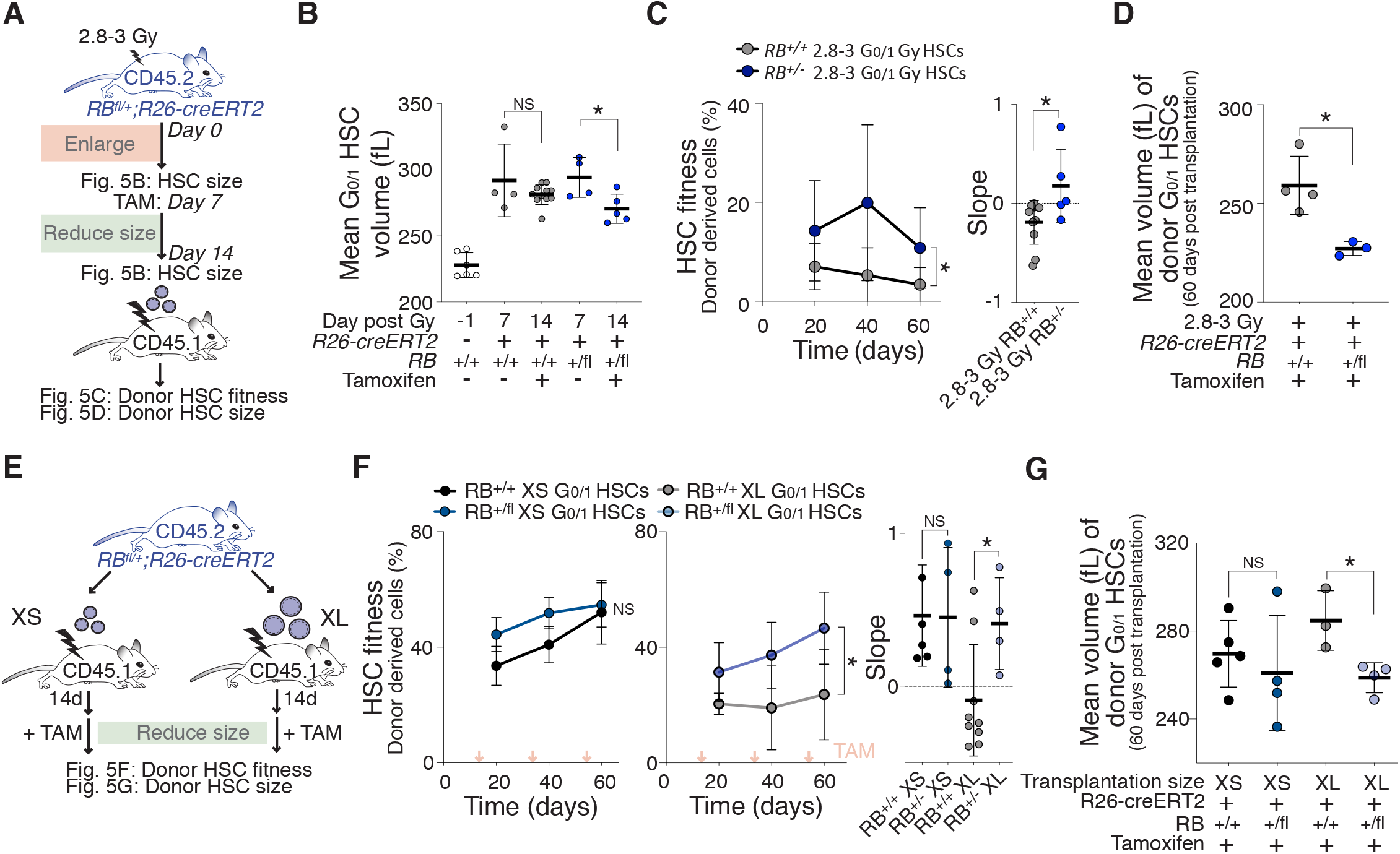
Reducing cellular size restores HSC fitness. (**A**) 8 week-old *R26-creERT2* mice carrying *RB*^+/+^ or *RB*^fl/+^ alleles were sub-lethally irradiated (2.8-3 Gy) and 7 days later HSC size was measured (B) and mice were treated with tamoxifen for 7 days. 14 days after irradiation, G_0/1_ HSC volume was measured (B) and 700 G_0/1_ HSCs were transplanted together with CD45.1 BM (420,000) into lethally irradiated recipient mice. Fitness (C) and volume (D) of donor G_0/1_ HSCs in recipient mice was measured. (**B**) Mean volume (fL) of *RB*^+/+^ or *RB*^fl/+^ *R26-creERT2* G_0/1_ HSCs isolated from mice 7 days after 2.8-3 Gy irradiation (*n* = 4) and 14 days after irradiation and treatment with tamoxifen for 7 days (*n* = 5-11) as described in (A) using a Coulter counter. Day −1 = before irradiation. Same control and 2.8-3 Gy sample as in Fig. 1A. (**C**) Percentage (%) of white blood cells derived from donor *RB*^+/+^ (*n;* donors = 6, recipients = 11) or *RB* ^fl/+^ (*n;* donors = 5, recipients = 5) *R26-creERT2* 2.8-3 Gy G_0/1_ HSCs that were treated before transplantation with tamoxifen as described in (A, 2 independent experiments). Slope of percentage (%) of donor-derived white blood cells in recipients over time. (**D**) Mean volume (fL) of recipient-derived donor *RB*^+/+^ (*n* = 4) or *RB* ^fl/+^ (*n* = 3) *R26-creERT2* 2.8-3 Gy G_0/1_ HSCs that were treated before transplantation with tamoxifen as described in (A) as measured by Coulter counter. (**E**) Schematic of reconstitution experiments: 600 XS- or XL-sized G_0/1_ HSCs were isolated from *R26-creERT2* mice carrying *RB*^+/+^ or *RB* ^fl/+^ alleles and transplanted together with CD45.1 BM (420,000) into lethally irradiated recipient mice. After transplantation, recipient mice were treated with tamoxifen at the indicated time points. Fitness (F) and volume (G) of donor G_0/1_ HSCs in recipient mice was measured. (**F**) Percentage (%) of donor HSC-derived white blood cells in recipients and slope of reconstitution were determined over time as described in (E)(*n;* donors = 4, recipients ≥ 4, 2 independent experiments). Red arrows indicate recipient treatment with tamoxifen. (**G**) Mean volume (fL) of G_0/1_ HSCs isolated from recipient mice that were reconstituted with donor XS- or XL-sized G_0/1_ HSCs from *R26-creERT2* mice carrying *RB*^+/+^ or *RB* ^fl/+^ alleles after they were treated with tamoxifen was measured by Coulter counter (*n* ≥ 3). Transplantation size displays the size of HSCs used for transplantation (XS or XL). Graph shows size of those HSCs 80 days after transplantation. For all panels, statistical significance was calculated using unpaired *t*-test to compare 2 samples, one-way ANOVA - multiple comparison - Tukey post-hoc test to compare multiple (3 or more) samples, ****P < 0.0001, ***P < 0.001, **P < 0.01, *P < 0.05; NS, not significant. Mean ± s.d is displayed.

Does reducing the size of naturally large HSCs also restore their reconstitution potential? To address this, we transplanted XL-sized *RB^fl/+^;R26-creERT2* HSCs. 14 days after transplantation, we treated animals with tamoxifen to excise one copy of the *RB* gene (**Fig. 5E**). This not only reduced the size of XL-HSCs, it also significantly increased the ability of XL-HSCs to build the hematopoietic compartment of recipients. *RB-/+* HSCs formed 47% of blood cells in recipients, whereas *RB+/+* HSCs contributed only 23% (**Fig. 5F, G**). Tamoxifen itself had no significant affect on HSC fitness (**fig. S8B**). Consistent with increased reconstitution potential, *RB* excised XL-HSCs more readily entered the cell cycle (**fig. S8C**). *RB* removal did not improve HSC fitness of small or control HSCs (**Fig. 5F**) (*46*), indicating that lack of Rb does not improve HSC function, but restores fitness to large HSCs. We conclude that the reduced fitness of naturally large HSCs can be improved by reducing their size. Our data demonstrate that HSC size is an important determinant of their *in vivo* fitness.

### HSC enlargement contributes to their functional decline during aging

Like many other stem cells, the function of HSCs declines with organismal age, with HSCs losing their ability to regenerate the hematopoietic system (*47*). Large HSCs share many of the characteristics of cellular aging: loss of stem cell potential, loss of proliferative abilities, up-regulated expression of genes associated with aging (**fig. S9A**), increased DNA-damage, and enlarged nucleoli (*48–50*). Based on these observations, we hypothesized that HSCs enlarge *in vivo* as mice age (*50*). We analyzed HSC size in mice that experience HSC decline at a younger age (D2 early-aging strain) and in the BL/6 strain, in which HSCs decline occurs later in life (*51*). HSCs were larger in old D2 (32 weeks), BL/6 middle-aged (56-65 weeks) and BL/6 old (86-102 weeks) animals compared to younger mice (8 weeks; **Fig. 6A-B**). This size difference was neither due to changes in cell cycle distribution (**Fig. S9B-C**), nor associated with changes in SA-β-gal activity (**fig. S9D**). As in response to irradiation, only a fraction of HSCs became enlarged, as judged by a broadening of the size distribution (D2 and BL/6, **fig. S9E**). One interpretation of this observation is that this change in cell size distribution is due to stochastic cellular damage that occurs naturally as some HSCs divide during aging while most other HSCs stay quiescent or do not experience damage during cell division (*52*).

**Fig. 6.**
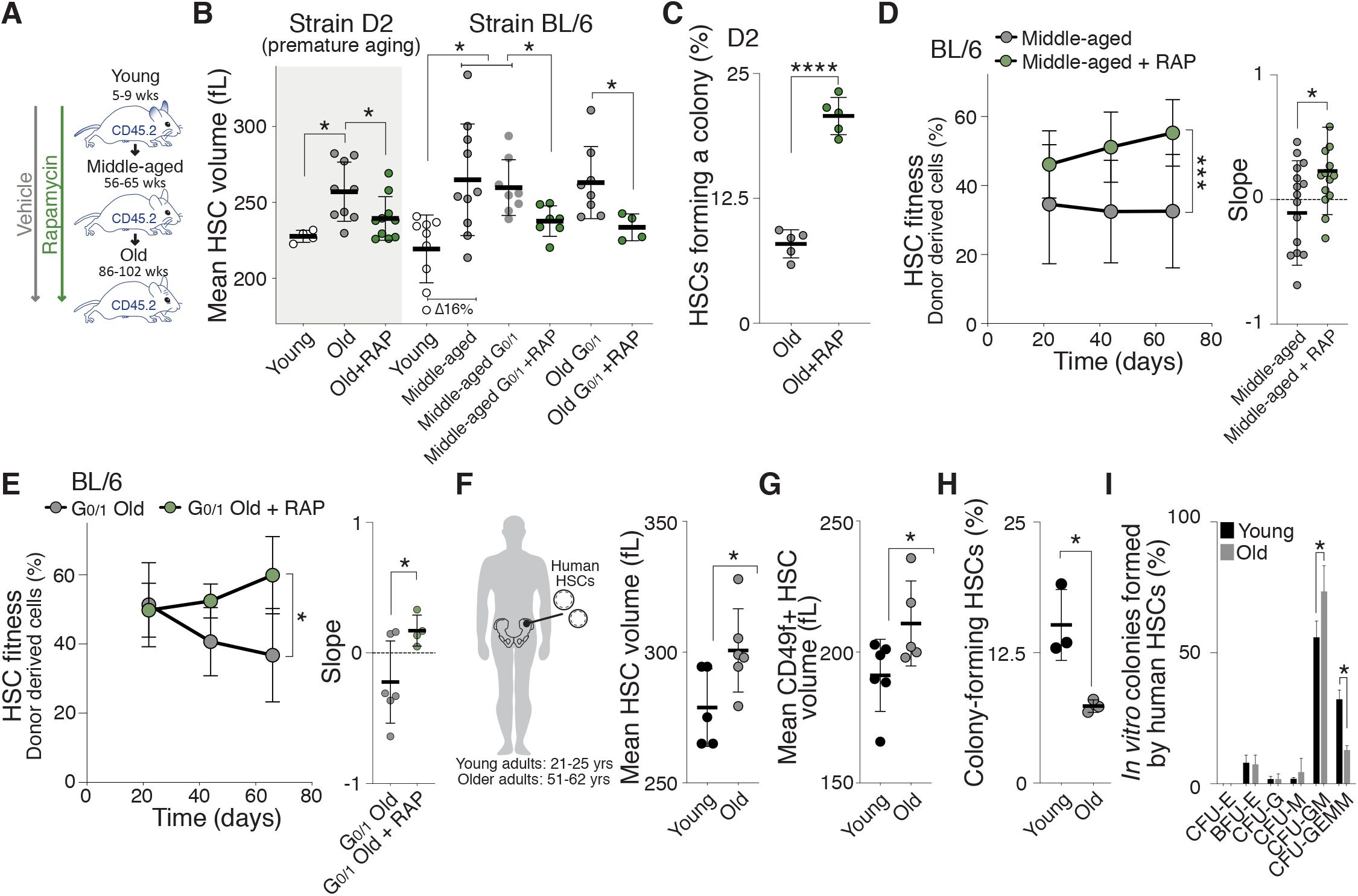
Enlargement of HSCs contributes to fitness decline during aging. (**A**) Mice were treated with vehicle or rapamycin during aging (from week 8 onward) and HSCs were isolated at 32 weeks (D2 mice) for volume analysis (B), *in vitro* colony forming assay (C) or at 56-65 weeks (middle-aged, BL/6) and 86-102 (old, BL/6) weeks for reconstitution assays (D-E). (**B**) Mean volume (fL) of HSCs from 7 week old (young, *n* = 4), 32 week old (old, *n* = 9) or old + rapamycin (RAP, *n* = 10) early-aging D2 mice and from 5-9 week old (young, *n* = 9), 56-65 week (middle-aged, *n* = 10), middle-aged G_0/1_ (*n* = 8), middle-aged + RAP (*n* = 7), 86-102 week old (old, *n* = 8) or old + RAP (*n* = 4) BL/6 mice was measured using a Coulter counter. Rapamycin treatment started at 8 weeks. Same WT BL/6 control as in Fig. 2H. (**C**) Colony forming efficiency *in vitro*: HSCs from D2 mice treated with vehicle or rapamycin during aging were plated on methylcellulose and percentage of HSCs forming a colony was quantified (*n* = 5). (**D**) Reconstitution assay: 800 donor (CD45.2)-derived HSCs from middle-aged (56-65 wks) BL/6 mice treated with vehicle (*n;* donors = 12, recipients = 16) or rapamycin (*n;* donors = 12, recipients = 14) during their life were transplanted together with CD45.1 BM (420,000) into lethally irradiated recipient mice (CD45.1). Percentage (%) of donor-derived white blood cells in recipients and slope of the reconstitution kinetics were measured over time (5 independent experiments). (**E**) Reconstitution assay: 1,000 donor (CD45.2)-derived G_0/1_ HSCs from old (86-102 wks) BL/6 mice treated with vehicle (*n;* donors = 6, recipients = 6) or rapamycin (*n;* donors = 4, recipients = 4) during their life were transplanted together with CD45.1 BM (420,000) into lethally irradiated recipient mice (CD45.1). Percentage (%) of donor-derived white blood cells in recipients and slope of the reconstitution kinetics were measured over time (2 independent experiments). (**F-G**) Mean volume (fL) of human HSCs (Lin-, CD34+, CD90+, CD38-, CD45RA- w/ and w/o CD49f+) from young (21-25 yrs, *n* ≥ 5) or old (51-62 yrs, *n* ≥ 5) individuals as measured by Coulter counter. (**H**) Colony forming efficiency *in vitro*: Human HSCs from young (21-25 yrs) or old (51-62 yrs) individuals were plated onto methylcellulose and the percentage of HSCs forming a colony was quantified after 21 days (*n* = 3). (**I**) Differentiation assay *in vitro*: Percentages of human GEMM, GM or single lineages (G, M, and E) colonies per plate were counted after 21 days (*n* = 3). For all panels, statistical significance was calculated using unpaired *t*-test to compare 2 samples, one-way ANOVA - multiple comparison - Tukey post-hoc test to compare multiple (3 or more) samples, ****P < 0.0001, ***P < 0.001, **P < 0.01, *P < 0.05; NS, not significant. Mean ± s.d is displayed.

To determine whether cellular enlargement contributed to the fitness decline of HSCs from aged mice, we prevented HSC enlargement during organismal aging. We treated young D2 and BL/6 mice from week 8 with rapamycin and found that aging-induced HSC enlargement was prevented (**Fig. 6A-B**). Importantly, these small-old D2 HSCs proliferated more and formed more colonies *in vitro* compared to large-old D2 HSCs (**Fig. 6C, S9F**). Rapamycin also preserved the reconstitution potential of old BL/6 HSCs. HSCs obtained from middle-aged or old BL/6 mice that were kept small by rapamycin treatment (**Fig. 6B**) were more effective in reconstituting lethally irradiated recipients than old-enlarged HSCs (**Fig. 6D-E, S9G**). Rapamycin had no effect on the fitness of HSC from young mice (**Fig. 1C, F)** or on *in vivo* differentiation (**fig. S9H**). These data indicate that HSC enlargement contributes to the fitness decline of HSCs during aging. However, rapamycin was not able to reverse HSC enlargement and age-induced loss of stem cell fitness. When we treated old animals (week 77 onwards) for 2-3 months with rapamycin, we did not observe a reduction in HSC size (**fig. S9I**) or recovery in HSC reconstitution potential (**fig. S9J**). This observation further indicates that rapamycin does not improve HSC fitness *per se*, but that its long-term effects on cell growth mediate preservation of HSC fitness. Our data suggest that HSC enlargement contributes to their functional decline during aging.

Lastly, we observed a similar relationship between cell size and aging in human HSCs (Lin^−^, CD34^+^, CD90^+^, CD38^−^, CD45RA^−^, ± CD49f^+^) (*53*). HSCs were larger in 51-62 year-old individuals compared to 21-25 year old individuals (**Fig. 6F-G**). Old-large human HSCs also exhibited a decreased ability to form colonies *in vitro* and to form multicellular lineages upon differentiation (**Fig. 6H-I**). We conclude that HSC enlargement during aging contributes to their functional decline in mice and humans.

## DISCUSSION

### Large cell size causes a decline in stem cell function

Stem cells are innately small in size. Here we show that size maintenance is important for HCS function. Conditions known to reduce stem cell potential – DNA damage and successive cell proliferation – cause an increase in HSC size. Two lines of evidence indicate that cell size affects HSC function: (i) Preventing HSC enlargement during insults such as DNA damage, aging and consecutive divisions protects HSCs from losing their stem cell potential; (ii) Reducing the size of large HSCs improves their fitness.

A key question regarding our findings is whether large HSCs are truly stem cells that have lost their stem cell potential or whether enlargement reflects a partially differentiated cellular state, in which multipotency had become restricted. We find that naturally and artificially enlarged HSCs are indistinguishable from smaller HSCs in their ability to home to the bone marrow niche and to differentiate into the various hematopoietic lineages *in vitro* and *in vivo*. The perhaps most compelling evidence that large HSCs are true stem cells is the observation that reducing the size of naturally large or irradiated HSCs improved their reconstitution potential.

It is important to note, that in response to DNA damage, proliferation challenge and aging, not all HSCs increase in size, but the size distribution of HSCs widens. This indicates that only a sub-population of HSCs enlarge during these challenges. We propose that this subpopulation is composed of HSCs that experienced DNA damage during irradiation and have divided more often during proliferation challenge and aging. A history of divisions increases the likelihood of HSCs to have experienced DNA damage that caused cell cycle arrest and hence cellular enlargement. Most other HSCs are quiescent. Their size will not be affected by insults such as DNA damage because mTOR activity is low during quiescence (*54*).

Our analysis of naturally large HSCs also revealed that they do not return to a smaller cell size when induced to proliferate during reconstitution assays. At first glance this result appears unanticipated. Several studies suggest that within their physiological size range, cells enter the cell cycle when they have reached a cell-type specific cell size, known as the critical size. When cells enlarge during cell cycle arrests the subsequent G_1_ is shortened and two smaller daughter cells are born (*55–61*). We hypothesize that this return to a smaller cell size only occurs when enlargement during cell cycle arrest does not exceed the critical size. When HSCs enlarge during prolonged cell cycle arrests beyond their critical cell size, restoration of the physiological size is no longer possible. In other words, they cannot shrink back to their natural size. Our data also indicate that HSCs do not grow in size indefinitely. Perhaps HSCs have an upper size limit, beyond which they die or faithful size homeostasis occurs at a different size set-point. However, it is clear from earlier studies that once cells become exceedingly large as occurs in cultured cells during senescence, multiple cellular processes, foremost cell division, fail (*16, 21, 22*). Our results suggest that the size window, in which HSCs function physiologically is small. Increasing HSC size by as little as 5-15 % leads to a decline in fitness.

### Causes of fitness decline in large HSCs

Our results indicate that *in vitro* proliferation capacity of HSCs is affected by large size. Given that cell proliferation is crucial for HSCs to rebuild the hematopoietic system of lethally irradiated mice, it is likely that reduced proliferative capacity is at least in part responsible for reduced fitness of large HSCs *in vivo*. Our previous work indicates that it is not the physical size of large cells that causes their reduced fitness, but rather the ensuing change in ratio of DNA to cytoplasm. Cellular enlargement in the absence of a corresponding increase in DNA content leads to a decrease in DNA:cytoplasm ratio. The concentration of unstable proteins decreases when this occurs in budding yeast (*16*). In naturally large HSCs, protein synthesis capacity and cell density of the whole cell are not changed, yet their fitness is decreased. Considering that HSCs have a very small cytoplasmic volume compared to their nuclear volume, the DNA:cytoplasm ratio might change dramatically even if overall size increases by only 20 %. As a result loss of unstable proteins specifically in the cytoplasm might contribute to impaired fitness upon enlargement. This is consistent with the observation that protein synthesis rates in HSCs must be tightly controlled (*62*). The observation that rapamycin treatment suppresses the fitness decline of HSCs during DNA damage further raises the possibility that mTOR-regulated processes besides cell growth, such as autophagy, or mitochondrial and lysosome function (*2*) influence HSC fitness during aging. Similarly, RB might improve the fitness of large HSCs by means other than cell size by acting on its other targets (*63*). Finally, our finding that large HSCs are depleted of metabolites like ATP might be an important clue for how enlargement causes dysfunction.

### A model for damage- and aging-induced loss of stem cell function

We have known since the 1960s that senescent mammalian cells are exceedingly large (*13, 14*). Similarly, aged yeast cells were first described to be large in the 1950s (*18*). These observations prompted us to investigate whether large cells harbor characteristics of senescence *in vivo*. We show here that DNA damage and aging lead to an increase in HSC size and that large size causes a decrease in HSC function. Young yeast and cultured primary human cells that are large also exhibit many of the phenotypes observed in senescent cells. Based on these observations, we propose that as stem cells divide and age, they experience DNA damage incurred during DNA replication or caused by telomere shortening (*34, 35*) (**Fig. S9K**). This DNA damage triggers cell cycle checkpoints and division transiently ceases to repair this DNA damage (*24, 36*). During these transient arrests, cell growth continues and HSCs increase in size (*15–17*). This enlargement then causes proliferation defects. In other words, it is not the division history itself that drives stem cell exhaustion, but the cellular enlargement resulting from past cellular damage incurred during cell divisions. As such, the mechanism of stem cell aging would be similar to replicative senescence observed in cultured human cells. We further propose that as large HSCs accumulate in the HSC population with age, smaller HSCs engage in more self-renewing divisions to compensate for the loss of functional stem cells (*52*), which might explain the observed upper limit of HSC size during aging. This model also provides an explanation for why very old mice harbor more cycling HSCs compared to young ones.

Our findings may have implications for rejuvenation therapies to improve stem cell function during aging. Rapamycin treatment has been reported to lengthen the lifespan of mice and improves the function of aged HSCs (*64, 65*). Our data indicate that rapamycin prevents the enlargement of HSCs and thereby protects them and potentially other stem cells from aging. However, rapamycin does not restore HSC function once they are large. We predict that compounds that revert cellular enlargement and hence reduce the size of already enlarged stem cells warrant further exploration as potential anti-aging therapies (*66*).

## Acknowledgements

We are grateful to J. Skotheim, J. Saarikangas, G. Neurohr, X. Zhou, D. Corbi, S. Morrill and the members of the Amon, Lees and Yilmaz lab for discussions and for critical reading of the manuscript. We thank Christy Chao, Charlie Whittaker, Dikshant Pradhan, the Flow Cytometry Core, the KI Genomics Core/MIT BioMicro Centre for analytical and technical support and S. Imada, J. Replogle, P. H. Hsu, K. Knouse and L. Zasadil for sharing material and protocols.

## Author contributions

J.L. and A.A. conceptualized the project, designed experiments and wrote the manuscript. J.L., M.B., C.C., P.M., M.M., E.S., K. M., C.R., J.D.S., C.W., J.K., T.M. and H.H. performed and analyzed experiments. S.J.M., J.A.L, L.A.B., A.A. and Ö.Y. provided reagents or advice on experiments. All authors helped editing the manuscript.

## Data and materials availability

All data are available in the main text or the supplementary materials. The accession number for the RNA-seq is GSE154335.

## Supplementary Materials

Materials and Methods

Figures S1-S9

Tables S1-S3

References (67-94)

## Materials and Methods

### Mice

All work was performed in accordance with the MIT Institutional Animal Care Facility and with relevant guidelines at Massachusetts Institute of Technology (protocol number: 0715-073-18, 0718-053-21).

Unless otherwise indicated, all experiments were performed with female C57BL/6J (BL/6) mice. 9-11 wks old B6.SJL-*Ptprc^a^Pepc^b^*/BoyJ (CD45.1, B6.SJL, JAX stock #002014) and 8-10 wks old C57BL/6J mice (CD45.2, JAX stock #000664) for reconstitution assays were purchased from The Jackson Laboratory (Bar Harbor, ME). Female DBA/2J (D2) mice (JAX stock #000671, Charles River stock #026) were used for aging studies as indicated. Lgr5-EGFP-IRES-CreERT2 mice (B6.129P2-Lgr5tm1(cre/ERT2)Cle/J, JAX stock #008875) were used for ISC isolation. The Ki67^RFP^ knock-in strain (Jax stock # 029802, *Mki67^tm1.1Cle^*/J) was used to determine cell cycle stages. *Rosa26CreERT2* mice (*67*) (B6.129-*Gt(ROSA) 26Sortm1 (cre/ERT2) Tyj*/J, JAX stock #008463) were crossed with *TSC1^fl/fl^* mice (*68*) (*Tsc1tm1Djk*/J, JAX stock #005680) to generate strain *TSC1^fl/fl^;R26-creERT2*. *RB*^fl/+^ (*Rb1^tm2Brn^*/J, JAX stock #026563) (*69*) were crossed with *Rosa26CreERT2* mice (B6.129-*Gt(ROSA) 26Sortm1 (cre/ERT2) Tyj*/J, JAX stock #008463) to generate *RB^fll+^; R26-creERT2* mice, which were used at week 8.

### HSC isolation and measurements

Murine bone marrow (BM)-derived HSCs were isolated as described previously (*70*). Briefly, BM was harvested by flushing the long bones. Red blood cells were lysed in ammonium-chloride-potassium (ACK, Thermo Fisher Scientific, A10492-01) buffer and samples were washed in Iscove’s Modified Dulbecco’s Medium (IMDM, Invitrogen, 21056-023) containing 2 % fetal bovine serum (FBS). Lineage positive cells were depleted using a mouse lineage cell depletion kit (Miltenyi Biotec, 130-090-858, 130-110-470) and the following antibodies from BioLegend CD150 (Cat#115913; RRID: AB_439796, Cat#115904; RRID: AB_313683), CD48 (Cat#103427; RRID: AB_10895922; Cat#103431; RRID: AB_2561462), Sca-1 (Cat#108114; RRID: AB_493596), CD117 (Cat#105812; RRID: AB_313221) and BD Biosciences CD150 (Cat#562651; RRID: AB_2737705; Cat#115911; RRID: AB_493599), CD48 (Cat# 103404; RRID: AB_313019), Sca-1 (Cat#562058; RRID: AB_10898185; Cat#108114; RRID: AB_493596) and CD117 (Cat#561074; RRID: AB_10563203) were used for staining. 7-AAD (Thermo Fisher Scientific, A1310) or propidium iodide (Sigma-Aldrich, 81845) was used for the identification of non-viable cells. Cells were sorted using an Aria cell sorter (Becton Dickinson).

Human BM samples were purchased from AllCells®. HSCs were purified as described previously (*71, 72*). Briefly, BM was diluted with Ficoll buffer (2 mM EDTA in phosphate-buffered saline (PBS)) at a ratio of 7:1. Mixtures were carefully layered over Ficoll-Paque Premium (Sigma, GE17-5442-02) at a ratio of 3:1 in a conical tube and centrifuged at 400 x g at 18°C for 35 min. The upper layer containing plasma and platelets was aspirated leaving the mononuclear cell layer undisturbed at the interphase. The mononuclear cell layer was mixed gently with Ficoll buffer and centrifuged at 300 x g at 18°C for 10 min. Cells were washed with 40 mL of Ficoll buffer and centrifuged at 200 x g at 18°C for 10 min. Cells were resuspended in 300 μL MACS buffer (2 mM EDTA, 5 mg/mL bovine serum albumin (BSA) in PBS) per 10^8^ cells. 100 μL of FcR Blocking Reagent (Miltenyi Biotec, 130-059-901) per 10^8^ cells and 100 μL of CD34 MicroBeads (Miltenyi Biotec, 130-046-702) per 10^8^ cells were added. For staining, cells were incubated with the following antibodies for 30 min at 4°C: CD38 (Cat#342371; RRID: AB_400453, BD Biosciences), lineage cocktail (CD3/14/16/19/20/56) [UCHT1, HCD14, 3G8, HIB19, 2H7, HCD56] (Cat#348803, BioLegend), CD90 (Thy1) (Cat#561969; RRID: AB_10895382, BD Biosciences), CD49f (Cat#551129; RRID: AB_394062, BD Biosciences) and CD45RA (Cat#560675; RRID: AB_1727498, BD Biosciences). HSCs were purified using a LS MACS Column (Miltenyi Biotec, 130-042-401) in a magnetic separator, resuspended in IMDM with 5 % FBS and sorted using an Aria cell sorter (Becton Dickinson).

### Cell cycle stage analysis

BM cells were resuspended at 10^6^ cells/mL in pre-warmed IMEM supplemented with 2 % FBS and 6.6 μg/mL Hoechst-33342 (Thermo Fisher Scientific, #H3570). After 45 min of incubation at 37°C in a water-bath, cells were washed with cold IMEM supplemented with 2 % FBS and kept at 4°C. Lineage positive cells were depleted and remaining cells stained using HSCs antibodies as described above. Because Hoechst-33342 can be toxic to cells, we also used the Vybrant™ DyeCycle™ Violet Stain (Thermo Fisher, #V35003) to stain DNA (5 μM, 10^6^ cells/mL) by incubating cells at room temperature (RT) for 30 min. Cells were not washed before cytometer analysis and sorting. To distinguish between G_0_ and G_1_ HSCs, we took advantage of Ki67-RFP mice and analyzed HSCs that were stained with Hoechst-33342 as described above using flow cytometry. For *in vivo* proliferation studies, 1.25 mg EdU was injected intraperitoneally into mice 24 h before sacrifice. EdU incorporation into the DNA of HSCs was evaluated according to the manufacturer’s instructions with the Click-iT® EdU Alexa Fluor® 488 Imaging Kit (Life Technologies, C10337).

### HSC volume measurements

Size of HSCs was determined using a Multisizer-3 Coulter Counter (Beckman Coulter) or microscopy. When using the forward scatter (FSC-A) of a flow cytometer to isolate HSCs of a specific size, the gates were set so that XS-HSCs encompass the 10 % smallest and XL-HSCs the 10 % largest cells in the HSC population. The M gate was set to isolate HSCs of mean size +/− 10 %. Cytometer-based HSC size fractionation was confirmed by measuring the absolute cell volume using the Coulter Counter. To determine HSC size using the Coulter Counter, HSC size distribution data from Coulter Counter measurements were fitted to a Gaussian distribution to calculate mean size and standard deviation (SD). If needed, bone marrow from test and control animals were frozen in FBS with 10 % DMSO at −80 °C and used for HSC isolation and size analysis later. Size evaluation by microscopy is described in the section entitled “Fluorescence assays”.

We also explored using flow cytometry (forward scatter) measurements to assess cell size, but this method was only able to detect large differences in cell size. Forward scatter (FSC-A) measurements are not an accurate method to determine size as several other parameters influence this measurement (wavelength of laser illumination, collection angle, refractive index of the particle, and flow medium). Thus, FSC-A values can differ between independent experiments making it difficult to compare experiments. We therefore determined the sensitivity and accuracy of the FSC-A size measurement by comparing them to size measurements by Coulter Counter. We calculated a ratio of the mean FSC-A value of experimental and control HSCs (**Materials and Methods fig. 1A-B**). A ration above 1 indicated an enlargement of the treated sample. Next, we compared this ratio to Coulter Counter measurements (**Materials and Methods fig. 1C**). The relationship between the ratio (mean FSC-A experimental/mean control FSC-A) and the percentage enlargement measured by Coulter Counter was not linear. Small increases in cell size where not detected by FSC-A measurement. This result indicates that FSC-A can only detect large differences in HSC volume.

### Reconstitution and homing assays

B6.SJL recipient mice (CD45.1) were irradiated with 9.5 Gy using a Cesium-137 irradiator (γ cell 40). Donor HSCs were transplanted 10 h post irradiation. HSCs were transferred under isoflurane anesthesia by retro-orbital injection using live HSCs (C57BL/6J, CD45.2) combined with 420,000 B6.SJL (CD45.1) supporting white blood cells in a volume of 100 μL per mouse. We chose this high number of HSCs in our transplantation experiments to obtain robust reconstitution even when utilizing HSCs with poor reconstitution capacity. For reconstitutions using HSCs from sub-lethally irradiated (3 or 5 Gy) or aged animals, we used 750-1000 HSCs because of their reduced potential to reconstitute lethally irradiated recipient mice.

Reconstituted animals were housed in sterile caging, administered antibiotic septra water and wet food on the floor of their cages and monitored twice daily for 14 days for activity, labored breathing/dyspnea and appearance. One half of the mice cages were placed on a heat mat with power control (QC supplies, 250220, 250230). Reconstitution ability of HSCs was determined 3, 6 and 9 weeks after reconstitution by peripheral blood analysis. Peripheral blood was collected in heparinized capillary tubes into sodium heparin (Sigma-Aldrich, H3149-100KU) diluted in PBS. Red blood cells were lysed in ACK buffer and washed in Hank’s Balanced Salt Solution (HBSS, Thermo Fisher Scientific, 14175-095) containing 2 % FBS. Cells were then incubated with antibodies according to the manufacturer’s specifications and analyzed with a BD FACSCelesta™ Flow Cytometer (BD Biosciences). Engraftment and lineage analysis was determined by staining peripheral blood with CD45.1 (BioLegend Cat#110708; RRID: AB_313497), CD45.2 (BioLegend Cat#109806;RRID: AB_313443), myeloid markers Gr-1 (BD Biosciences Cat#562709; RRID: AB_2737736), and CD11b (BD Biosciences Cat#562605; RRID: AB_11152949) or lymphoid markers B220 (BD Biosciences Cat#562922; RRID: AB_2737894) and CD3e (BD Biosciences Cat#562600; RRID: AB_11153670). The reconstitution potential of HSCs was further tested by performing secondary and tertiary transplants with 10^6^ BM obtained from primary or secondary recipients, respectively, 100 days after their reconstitution.

HSC fitness (donor contribution) was calculated by determining the fraction of CD45.2^+^ cells in total white blood cells (sum of CD45.2^+^ and CD45.1^+^ cells). Statistical significance was calculated using the values of the last time point of the reconstitution. Reconstitution kinetics was assessed by calculating the slope of the degree of donor contribution to the recipient’s blood over time (linear regression). We calculated the slope using the first time point and the last time point before the degree of donor contribution plateaued.

The percentage of myeloid and lymphoid cells was determined by quantifying the fraction of Gr-1^+^, CD11b^+^, B220^+^ or CD3e^+^ cells that were also CD45.2^+^. Each lineage marker fraction was normalized to a sum of 100 %.

Homing assays were performed as reconstitution experiments, except 6,000-8,000 donor HSCs were injected. The percentages of donor HSCs in the bone in relation to recipient BM-cells were determined 21 hours after reconstitution.

### RNA isolation

For RNA isolation, 10,000 XS, M, XL or not size sorted HSCs in G_0/1_ were isolated as described above but in the absence of FBS. HSCs were sorted into 750 μL TrizolLS (Thermo Fisher Scientific, 10296010). During the sort, the collection sample was vortexed every 10 min. Post-sort, the samples were brought to a total volume of 1 mL with DEPC treated water. Samples were stored at −80 °C or directly used for RNA extraction. To extract RNA, 200 μL chloroform was added to the sample and briefly vortexed at medium speed. After 2 min at RT, the sample was centrifuged at 12,000 x g for 15 min at 4 °C. 400 μL of the aqueous phase was removed and put into 1,400 μL RLT buffer (Qiagen) with 14 μL β-mercaptoethanol and mixed vigorously. 1 mL 100 % ethanol was added to each sample, mixed by pipetting, then loaded onto RNeasy Micro Kit (Qiagen, 74004) columns. The manufacturer’s directions were followed from the RW1 wash step and the sample was eluted into 14 μL DEPC water.

For RNA-seq analysis, samples were analyzed for RNA integrity using a Femto Analyzer. Then the Clontech v4 Low-Input RNA Kit was used for the PolyA library prep. RNA‐seq data were aligned to the mm10 mouse genome assembly and the ensemble version 88 annotation with STAR version 2.5.3a and gene expression was summarized with RSEM version 1.3.0 (*73*).

### RNA-seq analysis

The RNA-Seq data is available from the Gene Expression Omnibus under accession number GSE154335. Expression data were analyzed using 2 different data sets: (1) All human GO gene sets msigDb version 7.1 (*74*) and (2) mouse gene categories of HSCs based on previous studies (*75–79*) (**Table S2**).

Differential expression analysis was performed with R version 3.4.4 and DESeq2 version 1.16.1 (*80*). Differentially expressed genes were defined as those having an absolute log2 fold change greater than 1 and a false discovery rate (FDR) - adjusted p-value of less than 0.05 (*q*-value). Deseq2 and edgeR version 3.4.4 programs were used to perform principle component analysis. Data parsing and clustering was performed using Tibco Spotfire Analyst 7.6.1. Mouse genes were mapped to human orthologs using Pre-ranked. GSEA analysis was performed using javaGSEA version 2.3.0. Differential expression analysis was done by making comparisons between two biological replicates of L, M, and S cell sizes and un-fractioned pool of cells. Very few genes meet the differential expression thresholds of log2 fold change greater than 1 or less than −1 and an adjusted *p*-value less than 0.05 (**Table S3**). Pre-ranked GSEA (version 4.0.3) was run using the DESeq2 Wald statistic as ranking metric and human gene symbols with the MSigDB (https://www.gsea-msigdb.org/gsea/msigdb/index.jsp) of the C5 GO gene sets. Similar runs were performed with mouse gene symbols and a collection of manually curated mouse gene sets. The small number of differentially expressed genes in these comparisons limits the utility of these tools but the data produced are consistent with the GSEA results. To calculate the percentage of GO gene sets associated with a biological pathway that passed the filter of being different when comparing XS- or M-sized HSCs with XL-HSCs but not different when comparing XS with M-sized HSCs, we determined the total number of GO gene sets associated with the biological process and computed the percentage (**Table S1**).

### *In vitro* cultivation of murine and human HSCs

Flow cytometer-sorted murine or human HSCs were resuspended in 4 mL complete MethoCult GF M3434 medium or MethoCult™ H4435 Enriched (STEMCELL Technologies, 03434 and 04435) thawed at RT and 1 mL of the suspension was placed into 35 mm dishes in triplicates. Growth at 37°C in a 5 % CO_2_ chamber was monitored until colonies were visible (7d and 21d, respectively). Number of colonies and colony type were quantified manually.

### Measurements of single-cell density and volume using a suspended micro-channel resonator

HSCs were isolated as detailed above and cultured *ex vivo* for 2 days in expansion medium (StemSpan™ SFEM, #09600 supplemented with cytokines). Single-cell density and volume measurements were carried out using the suspended micro-channel resonator (SMR), as detailed previously (*81, 82*). Briefly, the SMR is a vibrating cantilever that has a fluidics micro-channel passing through it. As a cell is flown through the channel in the cantilever, the vibration frequency changes proportionally to the buoyant mass of the cell. The vibration of the cantilever is monitored using piezoresistors. On the other side of the cantilever, the cell is mixed into a denser fluid, in which the cell is flown back through the cantilever to obtain another measurement of the buoyant mass. As the density of the two fluids is known, the absolute volume and density of the cell can be calculated from the two buoyant mass measurements (*82*). This is then followed by flushing another cell into the system for a measurement. The HSC population was continuously sampled this way for 5h. The average density or volume of the cells did not change during the experiment. The two media, in which the cells were measured were expansion medium and expansion medium supplemented with 30% OptiPrep (Sigma-Aldrich, D1556-250ML). The SMR was maintained at RT throughout the measurements, while the sample of suspended stem cells were kept on ice. The stem cells were mixed by gentle pipetting approximately every 1h in order to avoid cell clumping. Cell doublets were removed during the data analysis using the node deviation signal of the SMR, which differentiates doublets from singlets (*83*). The geometry, physical dimensions, operation of the SMR and data analysis were identical to those reported in (*81*).

### Intestinal crypt isolation and organoid formation assay of ISCs

As previously reported (*84*) and briefly summarized here, small intestines were removed from animals, washed with cold PBS, opened longitudinally and then incubated on ice with PBS containing EDTA (10 mM) for 45 min. Tissues were then moved to PBS. Crypts were mechanically separated from the connective tissue by shaking or by scraping, and then filtered through a 70 μm mesh into a 50 mL conical tube to remove villus material and tissue fragments. Crypts were dissociated into single cells by trypsinization and sorted by flow-cytometry into crypt culture medium, a modified form of medium as described previously (*84*). Isolated ISCs and Paneth cells were then mixed (1:1 ratio, 2,000 cells each), centrifuged, resuspended in 5 μL crypt culture medium and then seeded onto MatrigelTM (Corning 356231 growth factor reduced, Corning, 356231) containing 1 μM JAG-1 protein (AnaSpec AS-61298, NC0243900) in a flat bottom 48-well plate (Corning 3548). The MatrigelTM and cells were allowed to solidify before adding 300 μL of crypt culture medium. Organoid-forming capacity of the sorted ISCs was quantified after 3-5 days of culture. Each frequency was quantified by n > 100 cells per each view and n = 3 views per sample. In average, each frequency was determined from 20 - 30 organoids/ 100 - 300 ISCs.

### Drug and irradiation treatments

PD (Palbociclib, PD-0332991, LC Laboratories; 30.4 mg/kg body weight) (*85*), rapamycin (LC Laboratories R-5000; 6.4 mg/kg body weight) (*86*) or vehicle was intraperitoneally injected at a volume of 100 μL every 48 h for the indicated treatment time. Average body weight of the mouse was estimated to be 25 g. Rapamycin was reconstituted in ethanol and PD in DMSO. Drugs were then diluted in 20 % DMSO, 40 % PEG 400 (JT Baker, U216-07) in PBS. Blood composition of treated animals was analyzed after 70 days of treatment. Hematocrit in the peripheral blood was determined. Due to the decline in the Body Condition Score of recipient mice (CD45.1) that received rapamycin treatment after reconstitutions, we reduced the dose frequency to every 72 h. Mice were treated from week 8 with rapamycin (6.4 mg/kg body weight) every 3 days until week 102. Animals were sub-lethally irradiated using a 3, 5 or 6 Gy dosing. HSCs were isolated 14 days after irradiation for cell size analysis, senescence assays and reconstitution assays. The Cre-recombinase was activated via tamoxifen i.p. injections of 2 mg/100 μL per dose. Injections were performed using a 26 G needle (BD Tuberculin Syringes, 309625).

### Fluorescence assays

For immunofluorescence analyses, Fisherbrand™ Superfrost™ Plus Microscope Slides were primed with 0.1 % polylysine for 5 min, washed with dH_2_O and air-dried. 15,000 sorted HSCs were distributed on slides and incubated for 1 h in a humidified chamber at RT. HSCs were fixed for 20 min at RT with freshly prepared 4% paraformaldehyde (PFA, pH 7.2) and then washed three times with PBS. HSCs were permeabilized for 20 min in 0.2 % Triton-X 100, washed three times with PBS, and blocked for 30 min using 10 % Donkey Serum (Sigma, D9663) in PBS. Cells were incubated with primary antibody in 10 % Donkey Serum in PBS overnight at 4 °C: anti-Tfam (1:200, Cat# ab131607; RRID: AB_11154693) and anti-P-Ser139 γH2XA (1:100, Cat# 2577; RRID: AB 218010). After HSCs were washed three times with PBS + 0.1 % Tween-20, the secondary antibody solution (1:500, goat anti-rabbit Alexa 488, Cell Signaling Technology, Cat#4412S; RRID: AB_1904025, 1:500 goat anti-mouse Alexa 488, Invitrogen, Cat# A-10680; RRID: AB_2534062) was added for 1 h at RT in the dark in 10 % Donkey Serum in PBS. Coverslips were mounted with ProLong Gold Antifade Reagent with DAPI (Invitrogen, Molecular Probes, P36935) and imaged after 12 h. Control slides were not treated with primary antibody. Images were acquired using a DeltaVision Elite microscope (Applied Precision) platform (GE Healthcare Bio-Sciences) equipped with a CoolSNAP HQ2 camera (Roper), 250W Xenon lamps, SoftWoRx software (Applied Precision). Deconvolution was performed using SoftWoRx software with default settings. Fluorescence quantification and cell diameter measurements were performed with the Fiji software package. Maximum intensity projections of 20 stacks with step size 0.7 μm images were analyzed with ImageJ. For fluorescence mean intensity measurement, a Region Of Interest (ROI) was drawn around the area of interest and the mean intensity was quantified. An identically sized ROI was put next to the area of interest to determine the background mean intensity. Background mean intensity was subtracted from the area of interest mean intensity to yield the mean intensity (AU). Cells that were 2.5 times larger than the mean were excluded from the analysis. When the distribution of mean intensity was similar between independent experiments, but the average mean intensity was different due to microscopy settings, the mean intensity of independent experiments was normalized among each other to the control sample.

To detect mitochondrial mass and ROS in HSCs, lineage-depleted BM cells were incubated at 37°C for 15 min with 50 nM of MitoTracker Green (Thermo Fisher Scientific, M7514) (*87, 88*) or 10 μM CM-H_2_DCFDA (Thermo Fisher Scientific, C6827) after the staining of HSC surface marker. To create a positive control for ROS, the BM was treated with 20 μM CCCP. The negative control was left unstained. Cells were analyzed using flow cytometry.

To detect senescent HSCs, we used C_12_FDG (#7188, Setareh Biotech) to measure the activity of the senescence biomarker SA-β-galactosidase in un-fixed HSCs by flow cytometry (*89*). For fluorescence detection of SA-β-galactosidase activity, 10^7^ BM cell/mL were incubated with 30 μM chloroquine (#C6628-25G, Sigma Aldrich) in IMDM supplemented with 2 % FBS at 37°C for 30 min. 32 μM C_12_-FDG, a β-galactosidase substrate that generates a fluorescent product upon cleavage, was added for 30 min in a 37°C water bath. Cells were washed with 2 % IMDM at 4°C, lineage positive cells were depleted, lin-negative cells stained with HSC antibodies as described above and analyzed by flow cytometry. Unchanged forward and side scatter measures of cells was asserted after dye-treatment. The gates for senescent cells were set based on unstained HSCs controls. Specificity of C_12_FDG detecting senescent cells was determined in comparison with results obtained with the senescence β-galactosidase staining kit (Cell Signaling Technology #9860) on IMR90 cells that were treated with 100 ng/mL doxorubicin (Sigma, #D1515-10MG) or untreated (fig. S1D). IMR90 cells (ATCC® CCL-186™) were cultured in DMEM (Invitrogen) supplemented with 10 % FBS, 2 mM L-glutamine and 100 U/mL penicillin/streptomycin at 37°C in a 5 % CO_2_ camber.

Intensity of the HSC cell surface markers CD48, Slamf1/CD150, cKit, Ly6a/Sca1 and CD34 was quantified using flow cytometry and represented as surface concentration (fluorescence intensity (AU)/μm^2^) using the cell surface area determined using a Coulter Counter.

### Imaging and quantification of fibrillarin-stained nucleolus

Immuno-fluorescent images were acquired using a laser-scanning confocal microscope (Leica TCS SP8-X) using the Lightning mode with a 100× oil immersion/1.46 NA objective using a 1.28 optical zoom. Leica Applications Suite X 3.5.5.19976 software was used for image acquisition. Images of 2328×2328 pixels were obtained using zoom factor 1.28, resulting in a pixel size of 39.18 ×339.18 nm and pixel dwell time of 344 nm. For imaging DAPI, a 405 nm UV laser line was used (10% intensity) and emission was collected at 441-479 nm using a hybrid detector. An argon laser was used for detection Alexa Fluor 488 (excitation 488 nm; 30% laser power; 2% intensity) and emission was collected at 510-531 nm using a hybrid detector. Sequential scanning was used to prevent crosstalk between the DAPI and the AlexaFluor 488 detection and gain was adjusted to prevent pixel saturation. A minimum of 300 cells was imaged using the Mark and Find module. Z-stacks were set individually for each location using a z-step size of 0.2.

The area of the nucleolus (marked by fibrillarin-AlexaFluor 488) and the area of the nucleus (marked by DAPI and acting as a proxy for cell size) were measured in Fiji (ImageJ 1.52p) for analysis. Each channel was subjected to Z-projection using the “standard deviation” method and converted to binary using default settings. Regions of interest (ROIs) were defined as being individual nucleoli in the fibrillarin-AlexaFluor 488 channel and as individual nuclei in the DAPI channel. The ROIs were measured and the overall nucleolar area (defined as the sum area of all nucleoli in the cell), as well as the number of nucleoli were related to nuclear size, which correlates to cell size (data not shown). The nuclear areas were grouped into three categories: XS (which included the smallest 10% of nuclei), M (which included the mean 20% of nuclei by size) and XL (which covered the largest 10% of nuclei).

### Comet Assay

DNA damage in freshly collected HSCs was assessed performing a Comet Assay. 15,000 HSCs were collected per sample.

#### CometChip fabrication and cell loading

The CometChip PDMS molds were fabricated using the protocols described by Wood et al. (2010). 1% UltraPure™ agarose (16500 Invitrogen) was dissolved in PBS and poured on top of a sheet of GelBond film (53761 Lonza). A mold with 25 μm diameter microposts was placed on the 1% UltraPure™ agarose. After 5 minutes, the mold was removed, leaving the agarose gel with microwells attached to the Gelbond film. The film was placed on a glass plate before being fixed between the plate and a bottomless 96-well plate (655000 Greiner BioOne). Once the gel was clamped in place, 50 μL of the HSC solution were pipetted into each of the 96-wells and captured in microwells by gravity. The gel was rinsed with PBS then covered with 1% UltraPure™ Low Melting Point Agarose (16520050 Thermo Fisher Scientific).

#### FPG Treatment and CometChip Assay

The CometChip gel was incubated overnight at 4 °C in freshly made alkaline lysis stock (2.5 M NaCl, 100 mM Na_2_EDTA, 10 mM Tris, 1 % Triton X-100) to allow for DNA unwinding. After overnight lysis, the CometChip was washed three times with PBS before being incubated for 1 h at RT in FPG (M0240S New England Biolabs) diluted 1:10^4^ in PBS. The CometChip was then incubated at 4 °C in alkaline electrophoresis buffer (0.3 M NaOH, 1 mM Na_2_EDTA) for 40 min at 4 °C. Next, electrophoresis was performed at 4°C in the alkaline electrophoresis buffer at 1 V/cm and a current of 300 mA. The CometChip was neutralized by two 15-min washes in neutralization buffer (0.4 M Tris-HCl at pH 7.5) at RT.

#### Fluorescence and CometChip Analysis

Following neutralization, the CometChip was stained with SYBR Gold (S11494, Invitrogen) at a concentration of 1:10^4^ diluted in PBS. Images were captured with the 4 x objective of a Nikon 80i upright microscope and analyzed using the Guicometanalyzer, a custom software written in MatLab (The Mathworks) (*90*).

### Metabolite measurements in HSCs using liquid chromatography-mass spectrometry (LC-MS)

#### Water Soluble Metabolites

Methods for the isolation of cells for metabolomics were previously described (*91*). Here in brief, during the isolation, cells were kept cold. FACSAria flow cytometer was washed with ethanol and Milli-Q deionized water before the experiment and HSCs were sorted with a sheath fluid of 0.5× PBS, prepared fresh using Milli-Q water (Millipore), and a 70-μm nozzle in a four-way purity sort mode. 10,000 HSCs were directly sorted into 150 uL acetonnitril:methanol:water (40:40:20) pre-chilled low-binding eppendorf tubes and maintained at 4 °C during sorting. No FBS was used during the procedure. Samples that were sorted for no cells were used as a control. After sorting, each sample was kept on dry ice for the duration of the experiment, and then stored at −80 °C. The supernatants were centrifuged at 16,000 x g for 20 min to remove any residual debris before analysis.

The LC-MS method involved hydrophilic interaction chromatography (HILIC) coupled to the Q Exactive PLUS mass spectrometer (Thermo Scientific). The LC separation was performed on a XBridge BEH Amide column (150 mm 3 2.1 mm, 2.5 mm particle size, Waters, Cat#176002889). Solvent A is 95%: 5% H2O: acetonitrile with 20 mM ammonium bicarbonate, and solvent B is acetonitrile. The gradient was 0 min, 85% B; 2 min, 85% B; 3 min, 80% B; 5 min, 80% B; 6 min, 75% B; 7 min, 75% B; 8 min, 70% B; 9 min, 70% B; 10 min, 50% B; 12 min, 50% B; 13 min, 25% B; 16 min, 25% B; 18 min, 0% B; 23 min, 0% B; 24 min, 85% B; 30 min, 85% B. Other LC parameters are: flow rate 150 ml/min, column temperature 25°C, injection volume 10 μL and autosampler temperature was 5°C. The mass spectrometer was operated in both negative and positive ion mode for the detection of metabolites (*92*). Other MS parameters are: resolution of 140,000 at m/z 200, automatic gain control (AGC) target at 3e6, maximum injection time of 30 ms and scan range of m/z 75-1000. Raw LC-MS data were converted to mzXML format using the command line “msconvert” utility (*93*). Data were analyzed via the El-Maven software (*94*).

### Statistical analysis and data availability

Each experiment was repeated with three mice or as indicated. Representative experiments are shown only if the experiment was repeated three times and the results of each one supported the same conclusion. Standard deviations (SD) of the mean of three independent data points are shown in graphs or as indicated. Asterisks indicate P values: ****P < 0.0001, ***P < 0.001, **P < 0.01, *P < 0.05; NS, not significant. For all panels, statistical significance was calculated using unpaired *t*-test to compare two samples, One Way Analysis Of Variance (ANOVA)-multiple comparison - Tukey post-hoc test to compare multiple (three or more) samples or otherwise specified. Values were only excluded if most extreme value in the data set was a significant outlier from the rest (P < 0.05) according to Grubbs’ test. No statistical method was used to predetermine sample size. All other data supporting the findings of the study are available from the corresponding authors upon request.

---

**Figure S1:**
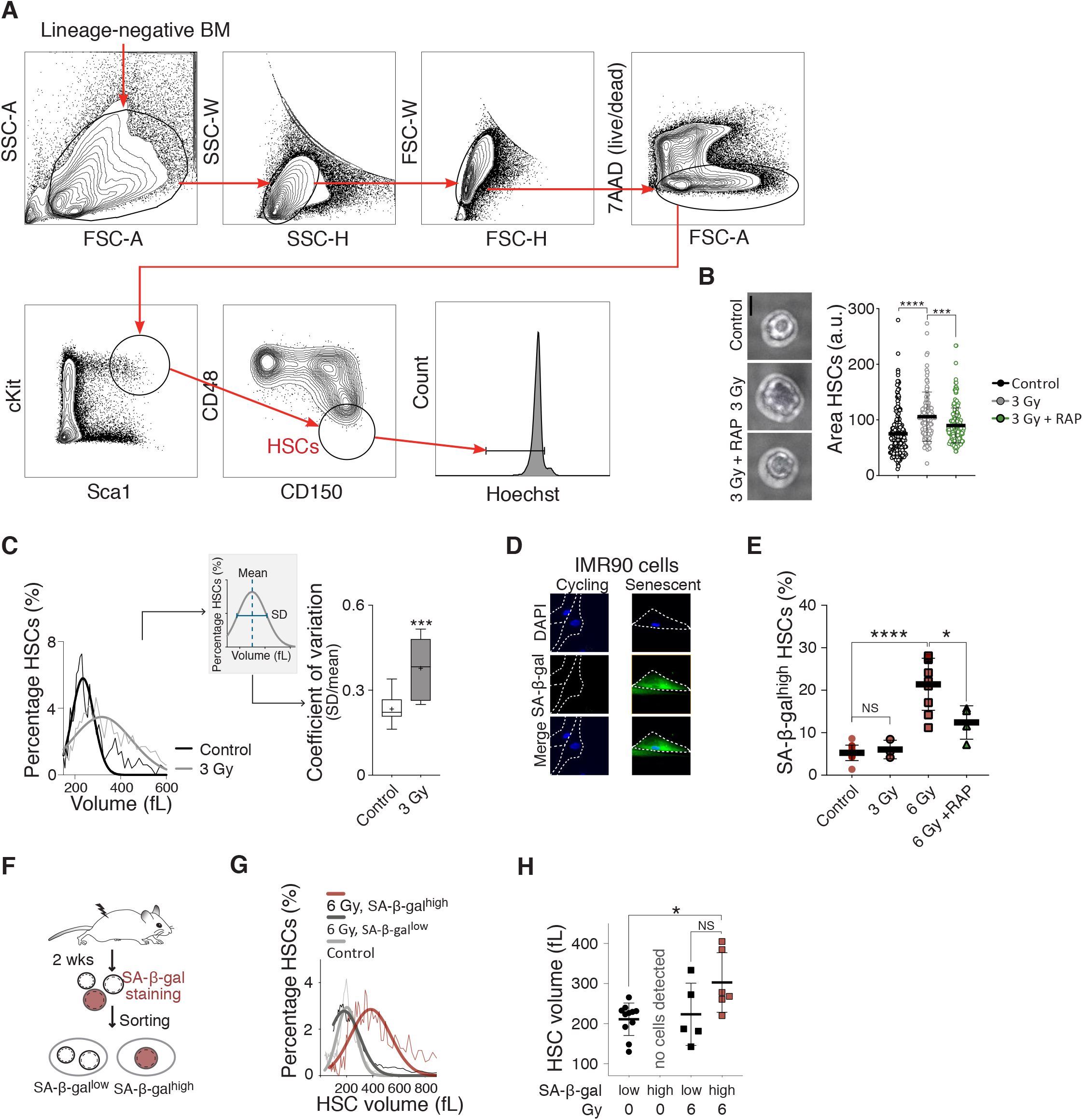
Enlargement of HSCs contributes to irradiation induced fitness decline. (**A**) Sorting protocol used to isolate BM-derived live G_0/1_ (Hoechst) HSCs (Lin-, Sca1/Ly6+, CD177/cKit+, CD150/Slamf1+, CD48/Slamf2-, 7ADD−) from an 8-12 week-old mouse in representative FACS plots. (**B**) Representative images of HSCs from control, sub-lethally irradiated (3 Gy) or 3 Gy + rapamycin (3 Gy + RAP) mice and quantification of HSC area (a.u., n = 2 independent experiments with a total of > 150 HSCs per condition analyzed, scale bar is 5 μm). (**C**) Representative raw data showing the size distribution: Percentage of HSCs (%) per volume (fL) from control or 3 Gy irradiated mice 2 weeks after treatment. Scheme of standard deviation (SD) and mean of Gaussian distribution fitted to the size distribution of HSCs to calculate coefficient of variation (CV). CV (SD/mean) of control HSCs (n = 15) or DNA-damaged HSCs (3 or 6 Gy, n = 6). (**D**) Positive control for SA-β-gal assay: Representative images of doxorubicin- or vehicle-treated primary human fibroblasts. SA-β-gal activity was assessed using the fluorescent substrate C_12_FDG. (**E**) Percentage of SA-β-gal^high^ HSCs (%) from control (n = 12), 3 Gy (n = 3), 6 Gy (n = 9) or 6 Gy + rapamycin (n = 4) mice 2 weeks after treatment. (**F**) Mice were irradiated with 6 Gy or left untreated. 2 weeks thereafter HSCs were sorted based on SA-β-gal staining intensity. (**G**) As described in (F): Representative raw data showing the size distribution of SA-β-gal^high^ and SA-β-gal^low^ HSCs from sub-lethally irradiated (6 Gy) and control mice. (**H**) As described in (F): Mean volume (fL) of SA-β-gal^high^ and SA-β-gal^low^ HSCs from 6 Gy (n ≥ 5) and control (n = 11) mice as determined by Coulter counter. For all panels, statistical significance was calculated using unpaired t-test to compare 2 samples, one-way ANOVA - multiple comparison - Tukey post-hoc test to compare multiple (3 or more) samples, ****P < 0.0001, ***P < 0.001, **P < 0.01, *P < 0.05; NS, not significant. Mean ± s.d is displayed.

**Fig. S2:**
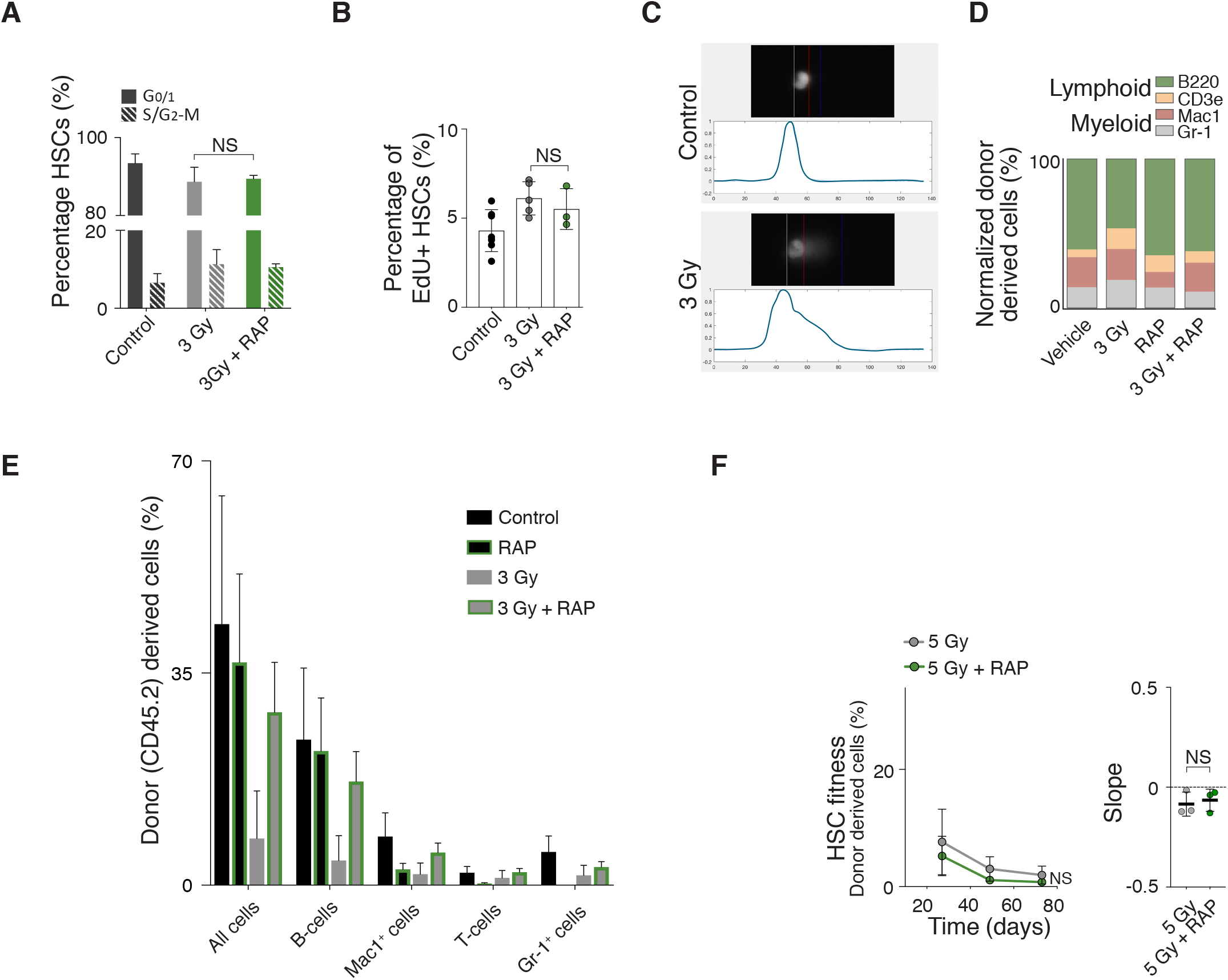
Enlargement of HSCs contributes to irradiation induced fitness decline. (**A**) Percentage of control (n = 3) and sub-lethally irradiated (3 Gy) HSCs treated with vehicle (n = 7) or rapamycin (n = 3) in the G_0/1_ and S/G_2_-M phases of the cell cycle as determined by DNA content (Hoechst 33342) analysis. (**B**) Percentage of control (n = 7) and sub-lethal irradiated (3 Gy) HSCs treated with vehicle (n = 5) or rapamycin (n = 3) that were positive for EdU incorporation. (**C**) Example of tail DNA measurement by CometChip assay to assess degree DNA damage. (**D**) Relative lineage distribution (lymphoid lineage B220, CD3e; myeloid lineage Mac1, Gr-1) of peripheral white blood cells from mice that were untreated or 3 Gy irradiated and afterwards treated with vehicle or rapamycin for 2 weeks (n = 4). (**E**) Reconstitution from Fig. 1C: Percentage of donor-derived total white blood cells (all CD45.2), B-cells (CD45.2 B220), T-cell (CD45.2 CD3e) and myeloid cells (CD45.2 Mac1, Gr-1) at day 60 post transplantation. (**F**) Donor (CD45.2) mice were treated with rapamycin or vehicle for 2 weeks, sub-lethally irradiated (5 Gy) and treated with rapamycin (RAP) or vehicle for another 2 weeks before CD45.2 HSCs were isolated and transplanted into lethally irradiated recipient mice with CD45.1 supporting BM (420,000) (n; donors = 3, recipients = 3). Recipient mice were not treated with rapamycin after the reconstitution. Percentage (%) of donor-derived white blood cells in recipients and slope of the reconstitution kinetics were measured over time. For all panels, statistical significance was calculated using unpaired t-test to compare 2 samples, one-way ANOVA - multiple comparison - Tukey post-hoc test to compare multiple (3 or more) samples, ****P < 0.0001, ***P < 0.001, **P < 0.01, *P < 0.05; NS, not significant. Mean ± s.d is displayed.

**Figure S3:**
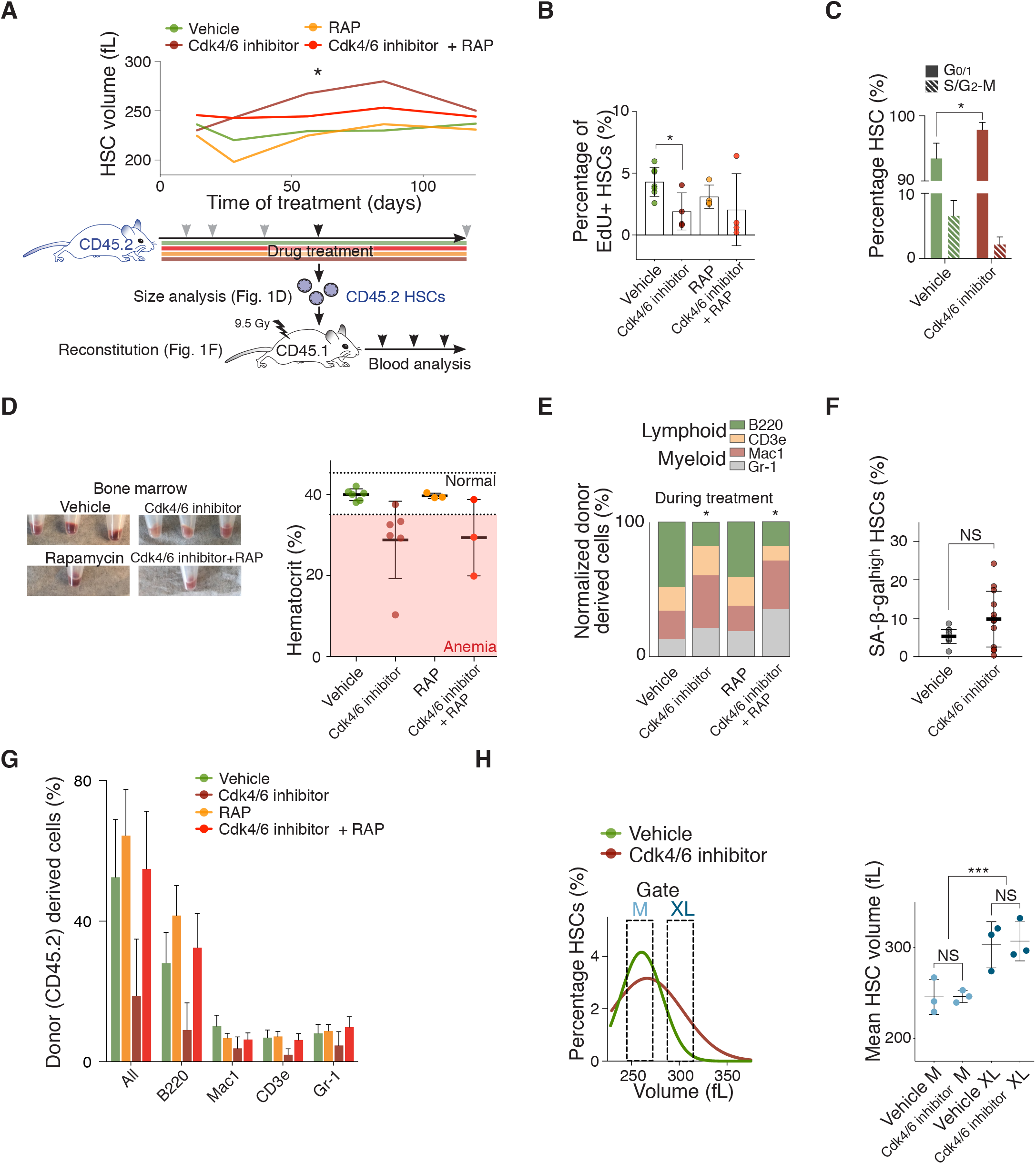
The Cdk4/6 inhibitor PD enlarges HSCs causing their decline in reconstitution potential. (**A**) Experimental design to artificially enlarge HSCs and subsequent analyses described in Fig. 1D-F. Mean volume (fL) of HSCs of vehicle, Cdk4/6 inhibitor (PD), rapamycin (RAP) or Cdk4/6 inhibitor + RAP treated mice was determined at the indicated time points using Coulter counter (grey and black arrows; total treatment time 165 days). (**B**) Percentage of HSCs from mice during the treatment with vehicle (n = 7), Cdk4/6 inhibitor (PD, n = 4), rapamycin (RAP, n = 4) or Cdk4/6 inhibitor + RAP (n = 4) that were positive for EdU incorporation. Same control as in Fig. S2B. (**C**) Representative images of bone marrow isolated from mice treated with vehicle, Cdk4/6 inhibitor (PD), rapamycin (RAP) or Cdk4/6 inhibitor + RAP for 85 days. The pale color of the bone marrow indicates anemia. Erythrocyte quantification represented as hematocrit of mice treated with vehicle (n = 6), Cdk4/6 inhibitor (PD, n = 6), rapamycin (RAP, n = 3) or PD + RAP (n = 3). Normal range (35.1 - 45.4 %); anemia (below 35.1 %). (**E**) *In vivo* differentiation assay: Relative lineage distribution (lymphoid lineage B220, CD3e; myeloid lineage Mac1, Gr-1) of peripheral donor-derived (CD45.2) white blood cells during 85 day treatment with vehicle (n = 4), Cdk4/6 inhibitor (PD, n = 4), rapamycin (RAP, n = 4) or Cdk4/6 inhibitor + RAP (n = 4). (**F**) Percentage of SA-β-galhigh HSCs (%) of mice treated with vehicle (n = 12) or Cdk4/6 inhibitor (PD, n = 13, Welch’s t-test). Same control as in Fig. S1E. (**G**) Reconstitution from Fig. 1F: Percentage of donor-derived total white blood cells (all CD45.2), B-cells (CD45.2 B220), T-cell (CD45.2 CD3e) and myeloid cells (CD45.2 Mac1, Gr-1) at day 60 post transplantation. (**H**) Size distribution of HSCs from mice treated with vehicle or Cdk4/6 inhibitor (PD) for 85 days. Gates used to isolate medium (M) and large (XL) HSCs are indicated. Mean volume (fL) of HSCs from mice treated with vehicle or Cdk4/6 inhibitor (PD) isolated using the M or XL gates was measured using a Coulter counter (n = 3). For all panels, statistical significance was calculated using unpaired t-test to compare 2 samples, one-way ANOVA - multiple comparison - Tukey post-hoc test to compare multiple (3 or more) samples or otherwise specified, ****P < 0.0001, ***P < 0.001, **P < 0.01, *P < 0.05; NS, not significant. Mean ± s.d is displayed.

**Figure S4:**
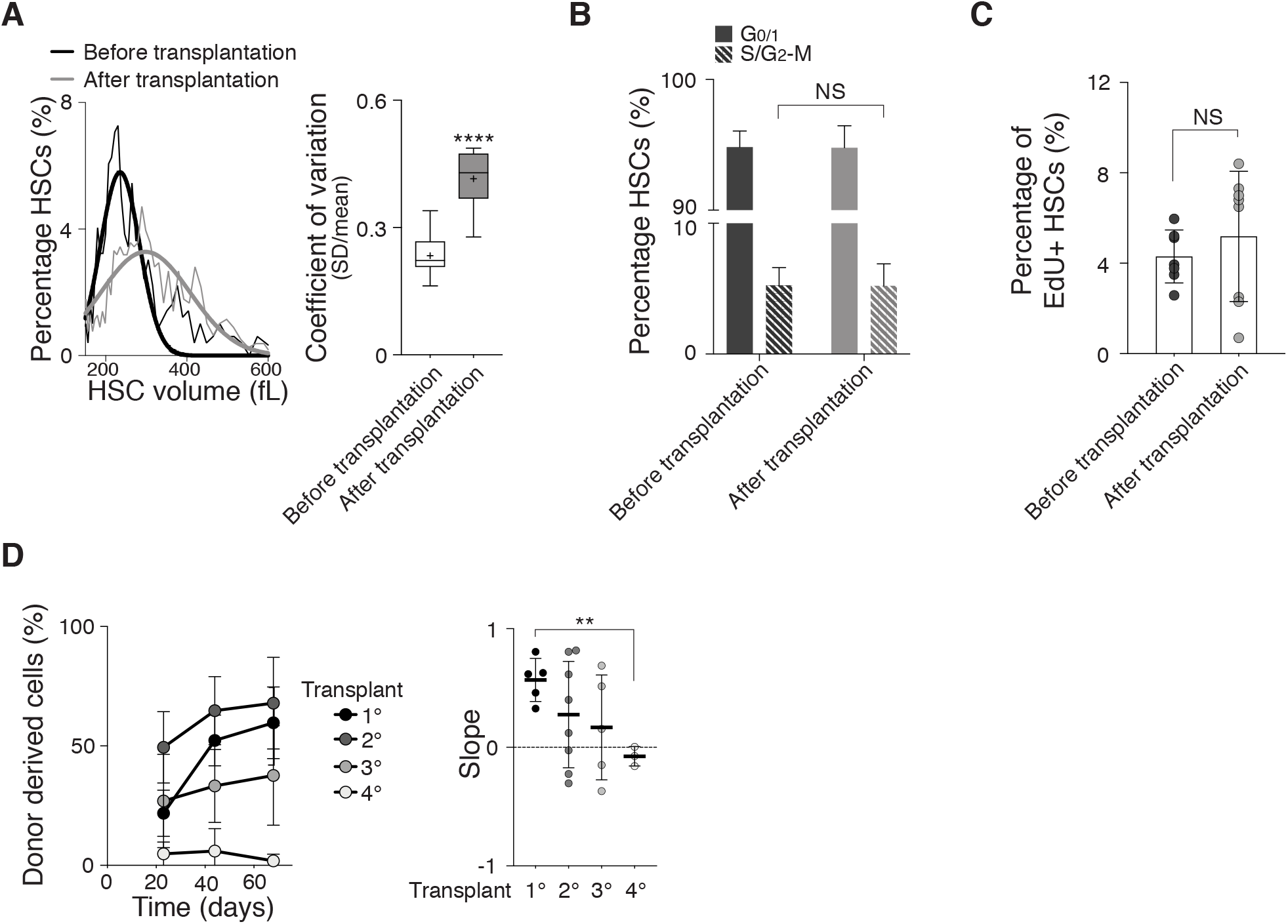
High cell division frequency enlarges HSCs contributing to their fitness decline. (**A**) Representative raw data showing HSC size distribution: Percentage of HSCs (%) per volume (fL) from before transplantation and after transplantation. Coefficient of variation (CV = SD/mean) of HSCs before transplantation (n = 15) or after transplantation (n = 9). Same control as in Fig. S1C. (**B, C**) Percentage of HSCs before and 80 days after reconstitution in the G_0/1_ and S/G_2_-M phases of the cell cycle as determined by (A) DNA content (Hoechst 33342, n ≥ 3) and (B) EdU incorporation analysis (n ≥ 7, same EdU control as in Fig. S2B). (**D**) Reconstitution assay: Donor HSCs were used in primary transplant and BM cells from previous transplants for following transplantations. Percentage (%) of donor-derived (CD45.2) white blood cells of primary (1°, n = 5), secondary (2°, n = 8), tertiary (3°, n = 5) and quaternary (4°, n = 3) transplants in recipients and slope of reconstitution kinetics over time. For all panels, statistical significance was calculated using unpaired t-test to compare 2 samples, one-way ANOVA - multiple comparison - Tukey post-hoc test to compare multiple (3 or more) samples, ****P < 0.0001, ***P < 0.001, **P < 0.01, *P < 0.05; NS, not significant. Mean ± s.d is displayed.

**Figure S5:**
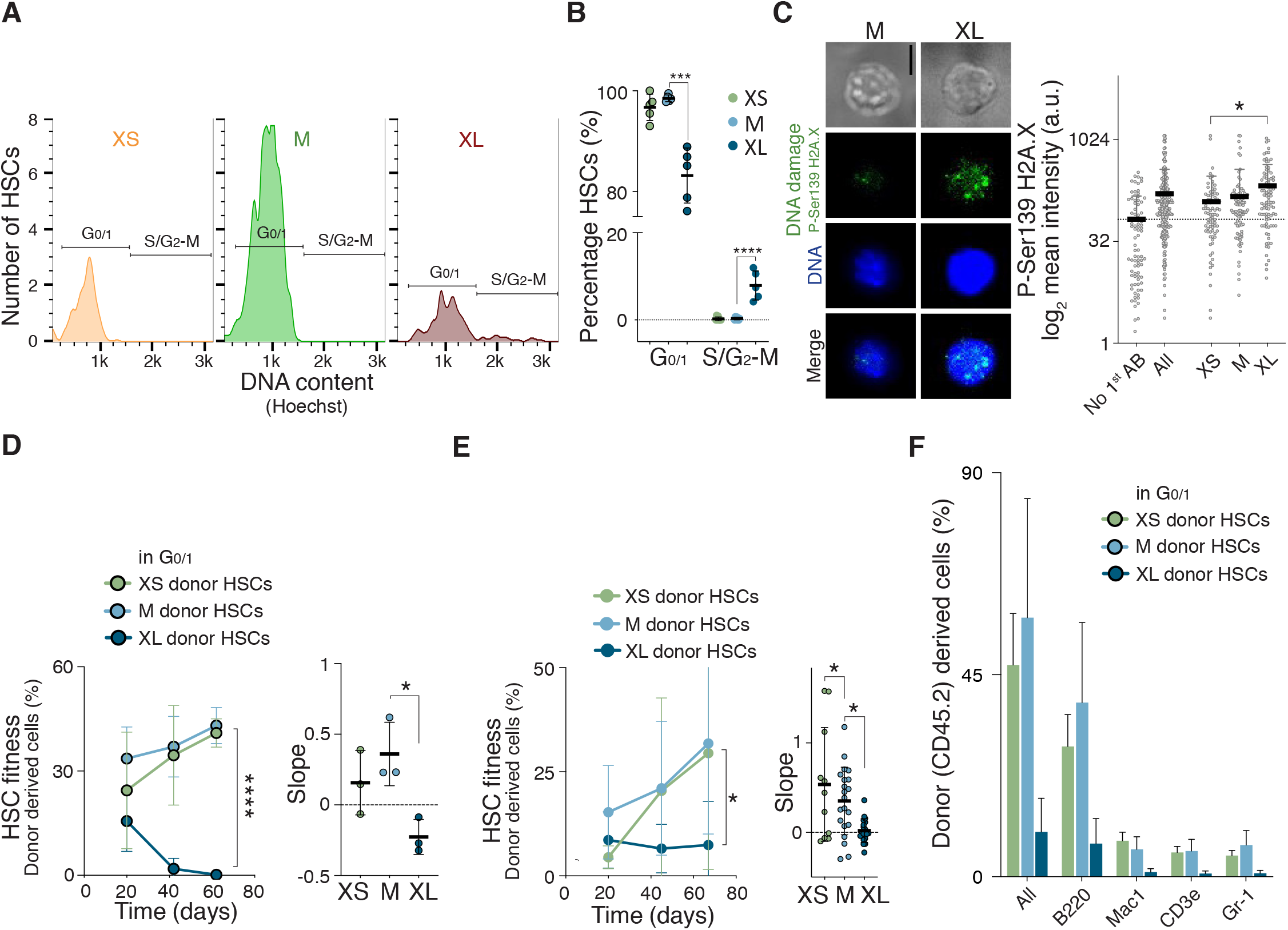
Naturally large HSCs are impaired in reconstituting the hematopoietic compartment. (**A**) Representative DNA content analysis of XS-, M- or XL-HSCs using Hoechst-33342. Gates of HSCs in G_0/1_ and S/G_2_-M are indicated. (**B**) Percentage of XS-, M- or XL-HSCs in the G_0/1_ and S/G_2_-M phases of the cell cycle as determined by DNA content (Hoechst) analysis (n ≥ 5). (**C**) Representative images of M- and XL-HSCs stained for P-Ser139 H2A.X (DNA damage) and DAPI (DNA), and quantification of fluorescence mean intensity (a.u., n = 3 independent experiments with a total of >100 cells per condition analyzed, scale bar = 5 μm). (**D**) Reconstitution assay: Percentage of donor-derived (CD45.2) white blood cells in recipient mice after transplantation of G_0/1_ Vybrant™ DyeCycle™ Violet (VD) stain-labelled donor XS-HSCs, M-HSCs or XL-HSCs together with CD45.1 BM (420,000) over time. Slope of percentage (%) of donor-derived white blood cells in recipient over time (n; donors = 2, recipients = 3). (**E**) Reconstitution assay: Percentage of donor-derived (CD45.2) white blood cells in recipients after transplantation of donor XS-HSCs (n; donors = 10, recipients = 11), M-HSCs (n; donors = 15, recipients = 21) or XL-HSCs (n; donors = 15, recipients = 19) together with CD45.1 BM (420,000) over time. HSCs were not stained with a DNA dye in this experiment. Slope of percentage (%) of donor-derived white blood cells in recipient over time (6 independent experiments). (**F**) Reconstitution from Fig. 3G: Percentage of donor-derived total white blood cells (all CD45.2), B-cells (CD45.2 B220), T-cell (CD45.2 CD3e) and myeloid cells (CD45.2 Mac1, Gr-1) at day 60 post transplantation. For all panels, statistical significance was calculated using unpaired t-test to compare 2 samples, one-way ANOVA - multiple comparison - Tukey post-hoc test to compare multiple (3 or more) samples, ****P < 0.0001, ***P < 0.001, **P < 0.01, *P < 0.05; NS, not significant. Mean ± s.d is displayed.

**Figure S6:**
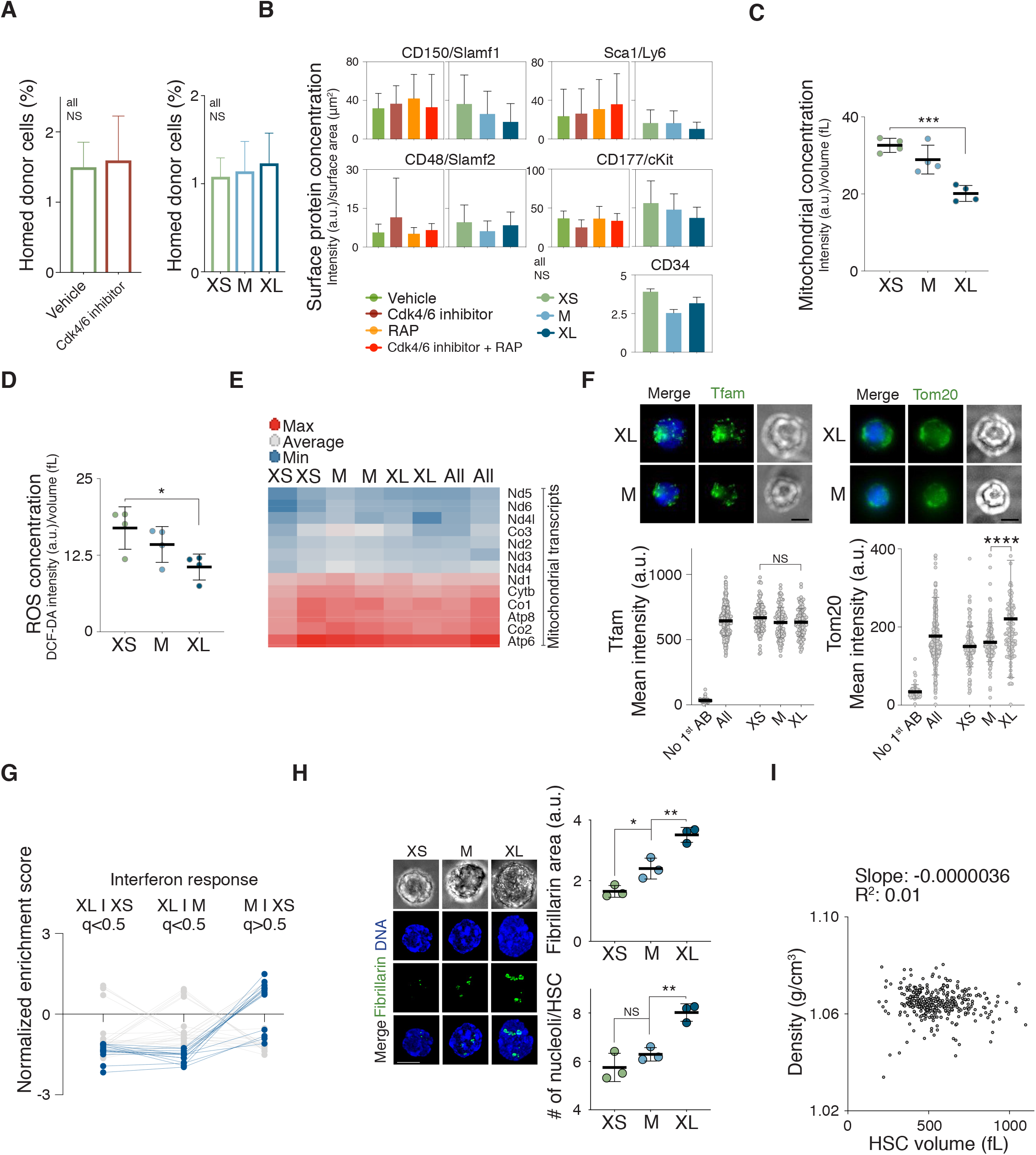
Characterization of large HSCs with DNA damage. (**A**) Homing assay: HSCs were isolated from mice treated with vehicle, Cdk4/6 inhibitor (PD), rapamycin (RAP) or Cdk4/6 inhibitor + RAP (n = 4) and XS-, M- or XL-HSCs from untreated WT mice (n = 2). The HSCs were transplanted into lethally irradiated recipient mice. Percentage (%) of donor-cells in the bone of recipient mice was measured 21 h after transplantation. (**B**) Fluorescence intensity (AU) of CD150/Slamf1, Sca1/Ly6, CD48/Slamf2, CD177/cKit and CD34 per surface area (μm2) was determined in differently sized HSCs or HSCs isolated from mice treated with vehicle, Cdk4/6 inhibitor (PD), rapamycin (RAP) or Cdk4/6 inhibitor + RAP (n ≥ 3). (**C**) Mitochondrial concentration (intensity (a.u.)/volume (fL) in XS, M and XL HSCs (n = 4). (**D**) ROS concentration (DCF-DA intensity (a.u.)/volume (fL) in XS, M and XL HSCs (n = 4). (**E**) Heat map of expression levels of mitochondrial genes in differently sized G0/1 HSCs (n = 2). (**F**) Representative images of mitochondrial protein Tfam and Tom20 in M- and XL-HSCs and DNA (DAPI), and quantification of fluorescence mean intensity (a.u., n = 2 independent experiments with each a total of > 110 cells per category analyzed, scale bar is 5 μm, AB = antibody). (**G**) Interferon response associated GO gene sets were analyzed using GSEA. Blue colored GO gene sets passed the filter to be different (q < 0.5) when comparing M- and XS-HSCs with XL-HSCs, but indistinguishable (q > 0.5) when comparing M-HSCs with XS-HSCs. Interferon response associated GO gene sets that did not pass the filter are displayed gray. (**H**) Representative images of XS-, M- and XL-sized HSCs from WT mice stained for fibrillarin and DNA (DAPI, scale bar is 5 μm). Quantification of total fibrillarin area (a.u.) and number (#) of nucleoli of XS-, M- and XL-sized HSCs (n = 3 independent experiments with a total of > 100 HSCs per condition analyzed). (**I**) HSCs were expanded in vitro for 2 days and the density (g/cm^3^) per HSC volume (fL) quantified (n = 351). For all panels, statistical significance was calculated using unpaired t-test to compare 2 samples, one-way ANOVA - multiple comparison - Tukey post-hoc test to compare multiple (3 or more) samples, ****P < 0.0001, ***P < 0.001, **P < 0.01, *P < 0.05; NS, not significant. Mean ± s.d is displayed.

**Figure S7:**
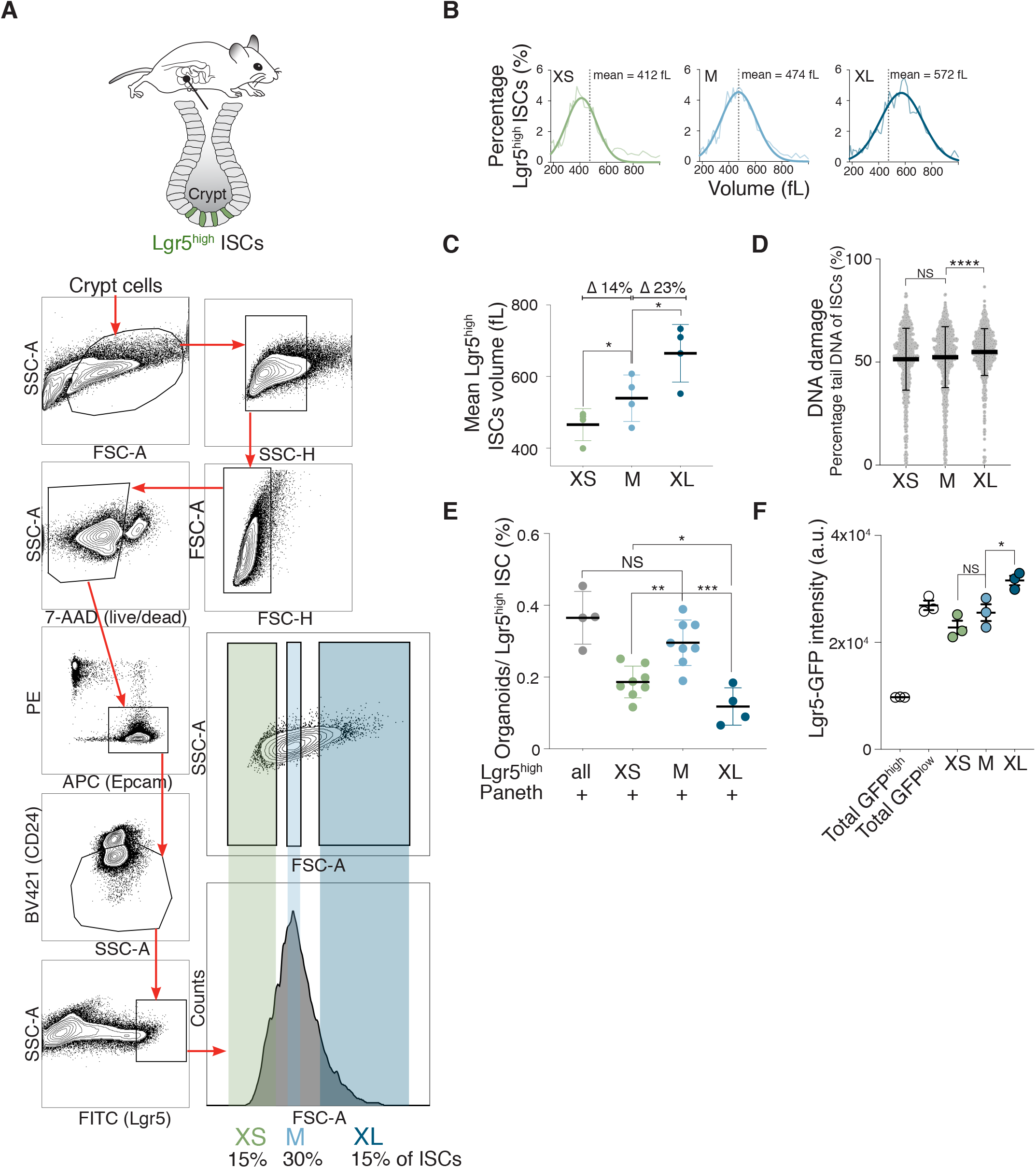
Size of intestinal stem cells affects their proliferation potential. (**A**) Schematic location of Lgr5high intestinal stem cells (ISCs) in the small intestine and sorting protocol used to isolate ISCs from an 8 week-old mouse. Representative FACS plots of cells of the crypt to enrich for ISCs (Epcam+, CD24-, Lgr5^high^) are shown. Histogram on the right shows the gates used to isolate small (XS), medium (M) and large (XL) ISCs. (**B**) Cell volume (fL) of XS-, M- or XL-ISCs was measured with a Coulter counter. Gaussian fit was used to determine mean cell volume. (**C**) Mean volume (fL) of ISCs isolated using the XS, M or XL gates as measured by Coulter counter (n = 4, Δ = difference). (**D**) CometChip assay to measure DNA damage: Percentage tail DNA of XS-, M- and XL-sized ISCs (n ≥ 729). (**E**) Percentage of organoids formed by all (n = 4), XS-(n = 8), M-(n = 8) or XL-(n = 4) Lgr5high ISCs co-cultured with Paneth cells. (**F**) Lgr5-GFP intensity (a.u.) of XS-, M- and XL-ISCs and total ISCs with high and low GFP intensity (n = 3). For all panels, statistical significance was calculated using unpaired t-test to compare 2 samples, one-way ANOVA - multiple comparison - Tukey post-hoc test to compare multiple (3 or more) samples, ****P < 0.0001, ***P < 0.001, **P < 0.01, *P < 0.05; NS, not significant. Mean ± s.d is displayed.

**Figure S8:**
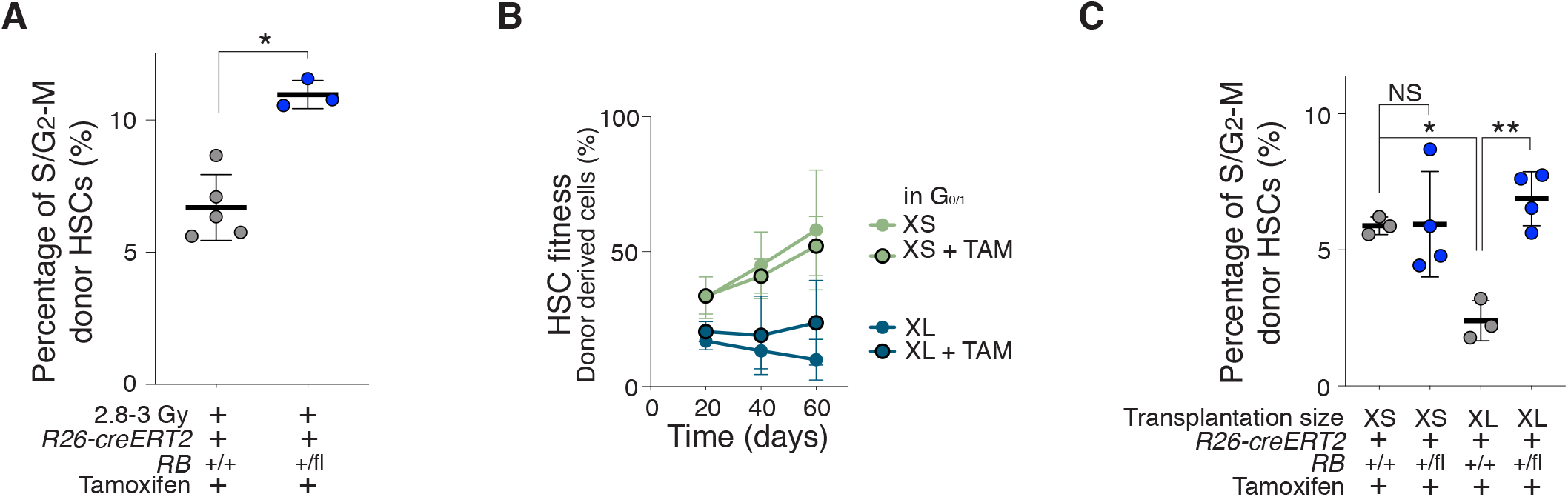
Reducing the size of large HSCs restores their reconstitution potential. (**A**) Percentage of 2.8 Gy irradiated *RB*^+/+^ (n = 5) or *RB*^+/−^ (n = 3) HSCs that are in S/G_2_-M from reconstituted animals as determined by DNA content (Hoechst 33342) analysis 60 days after reconstitution. (**B**) 600 differently sized donor-derived HSCs together with CD45.1 supporting BM cells (420,000) were transplanted into lethally irradiated recipient mice. Percentage (%) of donor-derived white blood cells in recipients after transplantation of G_0/1_ XS-HSCs (n; donors = 5, recipients = 6), or XL-HSCs (; donors = 5, recipients n = 6) treated with or without tamoxifen over time. Same data as in Fig. 3G and 5F. (**C**) Percentage of donor *RB*^+/+^ XS-(n = 3), *RB*^+/−^ XS-(n = 4), *RB*^+/+^ XL-(n = 3), *RB*^+/−^ XL- (n = 4) HSCs that are in S/G_2_-M from reconstituted animals as determined by DNA content (Hoechst 33342) analysis 60 days after reconstitution. For all panels, statistical significance was calculated using unpaired t-test to compare 2 samples, one-way ANOVA - multiple comparison - Tukey post-hoc test to compare multiple (3 or more) samples, ****P < 0.0001, ***P < 0.001, **P < 0.01, *P < 0.05; NS, not significant. Mean ± s.d is displayed.

**Figure S9:**
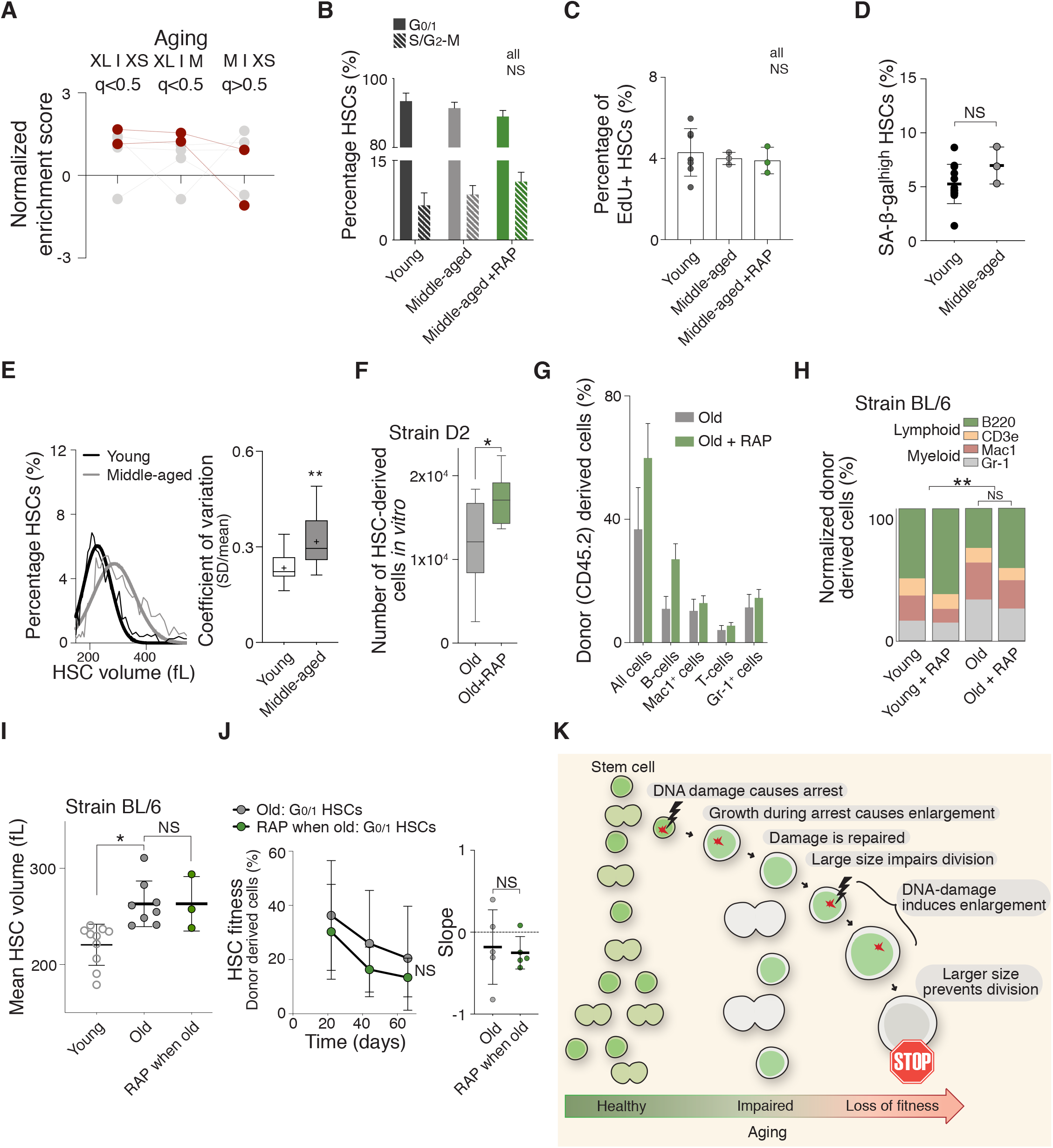
HSC enlargement contributes to their functional decline during aging. (**A**) Aging associated GO gene sets were analyzed using GSEA. Red colored GO gene sets passed the filter to be different (q < 0.5) when comparing M- and XS-HSCs with XL-HSCs, but indistinguishable (q > 0.5) when comparing M-HSCs with XS-HSCs. Aging associated GO gene sets that did not pass the filter are displayed gray. (**B, C**) HSCs from young (5-9 wks, n = 3-7), middle-aged (56-65 wks, n = 3) or middle-aged + rapamycin (56-65 wks, n = 3) BL/6 mice in the G_0/1_ and S/G_2_-M phase of the cell cycle as determined by DNA (Hoechst) content analysis (F) or EdU incorporation (G). Same control as in Fig. S2A (Hoechst) and S2B (EdU). (**D**) Percentage of SA-β-gal^high^ HSCs (%) from young (n = 12) and old (n = 3) mice. Same control as in Fig. S1E. (**E**) Representative raw data showing the size distribution: Percentage of HSCs (%) per volume (fL) from young or middle-aged mice. Coefficient of variation (CV = SD/mean) of HSCs from young (n = 15) or middle-aged (n = 10) BL/6 mice. Same control as in Fig. S1C. (**F**) Colony forming efficiency in vitro: HSCs from D2 mice treated with vehicle or rapamycin during aging were plated on methylcellulose and cell number was quantified (E, n = 9). (**G**) Reconstitution from Fig. 6E: Percentage of donor-derived total white blood cells (all CD45.2), B-cells (CD45.2 B220), T-cell (CD45.2 CD3e) and myeloid cells (CD45.2 Mac1, Gr-1) at day 60 post transplantation. (**H**) *In vivo* differentiation assay: Relative lineage distribution (lymphoid lineage B220, CD3e; myeloid lineage Mac1, Gr-1) from peripheral donor-derived (CD45.2) white blood cells 80 days after recipient mice were reconstituted with HSCs from 8 (young) or 102 (old) week-old BL6 mice treated with vehicle or RAP. No drug treatment was performed after the reconstitution (n = 4). (**I**) Mean volume (fL) of HSCs from 5-9 week old (young, n = 9), 86-102 week old (G_0/1_ old, n = 8) or old mice treated with rapamycin for 2-3 month starting at week 77 (G_0/1_ RAP, n = 3) BL/6 mice was measured using a Coulter counter. Same young and old data as in Fig. 6B. (**J**) Reconstitution assay: Lethally irradiated recipient mice (CD45.1) were reconstituted with CD45.1 BM cells (420,000) and donor (CD45.2)-derived G_0/1_ HSCs from old (86 wks) BL/6 mice treated with rapamycin (n; donors = 5, recipients = 5) for 2-3 months during old age (week 77 onwards) or untreated (n; donors = 5, recipients = 5). Percentage (%) of donor-derived white blood cells and slope of the reconstitution kinetics over time in recipients (**K**) Model for how HSC enlargement contributes to their functional decline with age. With age, HSCs are more likely to have encountered stochastic cellular damage during divisions. This DNA damage causes a transient cell cycle arrests, during which mTOR continues to promote cellular growth and enlarges HSCs. This enlargement reduces the ability of HSCs to build a blood system. For all panels, statistical significance was calculated using unpaired t-test to compare 2 samples, one-way ANOVA - multiple comparison - Tukey post-hoc test to compare multiple (3 or more) samples, ****P < 0.0001, ***P < 0.001, **P < 0.01, *P < 0.05; NS, not significant. Mean ± s.d is displayed.

**Experimental procedures Fig. 1.**
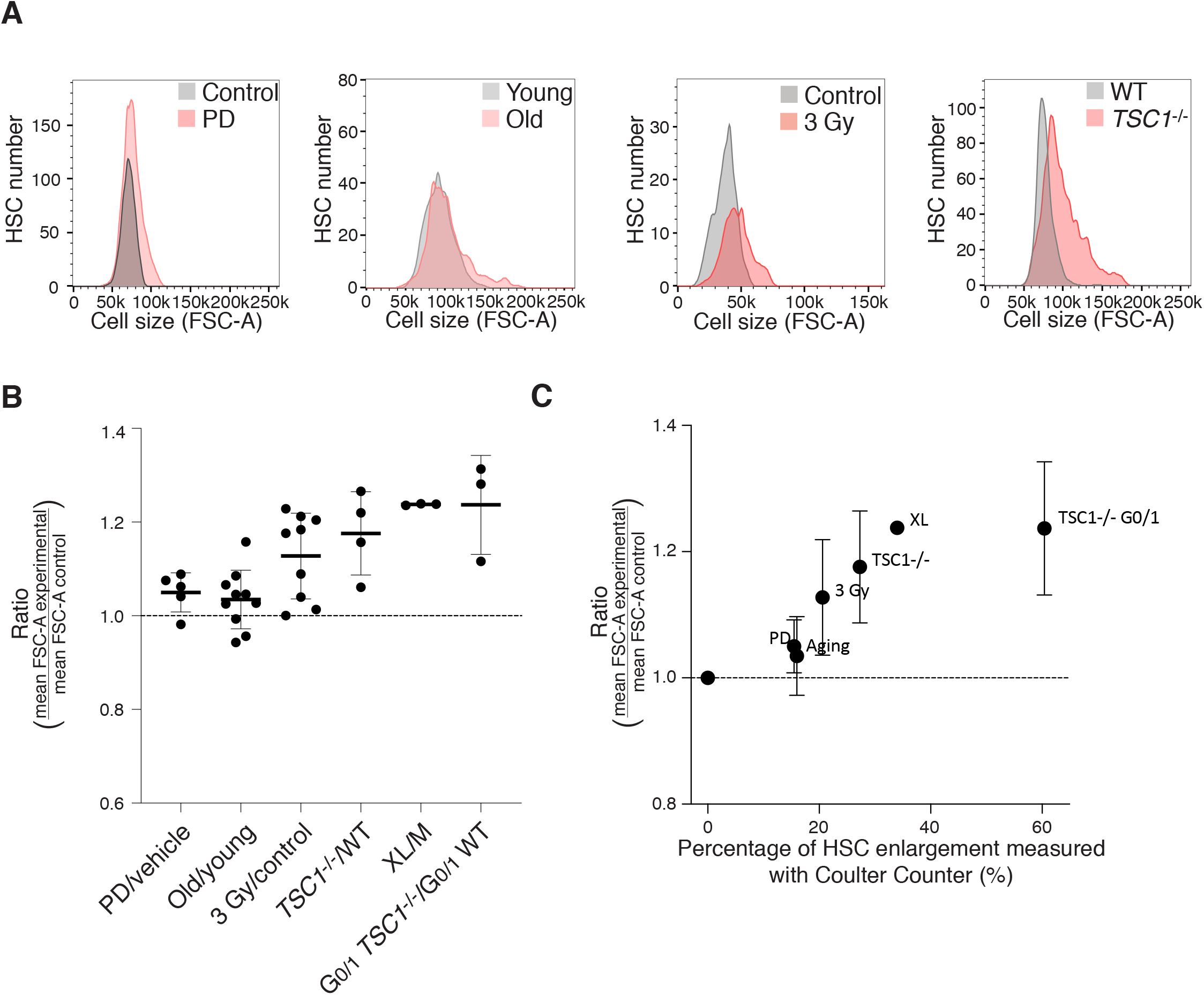
Flow cytometry (forward scatter) measurements only detect large differences in HSC size. (**A**) Histogram of forwards scatter (FSC-A) of HSCs enlarged by different methods. HSCs were compared to controls isolated in the same independent experiment. (**B**) Ratio of mean forward scatter (FSC-A) value of experimental HSCs and control HSCs. PD/vehicle (n = 5), old/young (*n* = 10), 3 Gy/control (*n* = 9), *TSC1*^−/−^/WT (*n* = 4), XL/M (*n* = 3), G_0/1_ *TSC1*^−/−^/G_0/1_ WT (*n* = 3). All measurements are from G_0/1_ HSCs, except for *TSC1*^−/−^ and PD treatment, for which all HSCs were analyzed (mean ± s.d). (**C**) Comparison of ratios measured in (B) to percentage of HSC enlargement measured by Coulter Counter (mean ± s.d). See also section” HSC volume measurements” in Materials and Methods.

